# Identification of potential inhibitory analogs of metastasis tumor antigens (MTAs) using bioactive compounds: revealing therapeutic option to prevent malignancy

**DOI:** 10.1101/2020.10.19.345975

**Authors:** Anik Banik, Sheikh Rashel Ahmed, Emran Hossain Sajib, Anamika Deb, Shiuly Sinha, Kazi Faizul Azim

## Abstract

The deeper understanding of metastasis phenomenon and detection of drug targets could be a potential approach to minimize cancer mortality. In this study, attempts were taken to unmask novel therapeutics to prevent metastasis and cancer progression. Initially, we explored the physiochemical, structural and functional insights of three metastasis tumor antigens (MTAs) and evaluated some plant based bioactive compounds as potent MTA inhibitors. From 50 plant metabolites screened, isoflavone, gingerol, citronellal and asiatic acid showed maximum binding affinity with all three MTA proteins. The ADME analysis detected no undesirable toxicity that could reduce the drug likeness properties of top plant metabolites. Moreover, molecular dynamics studies revealed that the complexes were stable and showed minimum fluctuation at molecular level. We further performed ligand based virtual screening to identify similar drug molecules using a large collection of 3,76,342 compounds from DrugBank. The results suggested that several structural analogs (e.g. Tramadol, Nabumetone, DGLA, Hydrocortisone) may act as agonist to block the MTA proteins and inhibit cancer progression at early stage. The study could be useful to develop effective medications against cancer metastasis in future. Due to encouraging results, we highly recommend further *in vitro* and *in vivo* trials for the experimental validation of the findings.

## 1. Introduction

Diseases are clinical conditions which impairs normal functions and structures of an organism [1]. However, among various types of morbid diseases, cancer is the most dangerous one [2, 3]. Cancer is a disease that can be found in almost any tissue or organ when abnormal cells uncontrollably form. Gradually, tumor cells exceed their natural boundaries, invade neighboring body regions and become malignant. Millions of people have to die every year due to the severity of this disease and the disease becomes more severe when metastasis occurs [4]. The International Cancer Research Agency focused on the geographical variation across 20 regions of the world and estimated that there will be 18.1 million new cancer cases and 9.6 million cancer deaths in 2018 [5]. Lung cancer is the most frequently diagnosed cancer in both sexes (11.6% of total cases) and the leading cause of cancer mortality (18.4%), followed by female breast cancer (11.6%), prostate cancer (7.1%), and colorectal cancer (6.1%) for prevalence and colorectal cancer (9.2%), stomach cancer (8.2%), and liver cancer (8.2%) for mortality [5]. Approximately 1,500 people continue to die from cancer each day, providing evidence for the inability to treat the disease as it spreads across the body [6].

Metastasis refers to the process by which the cancer cells migrate through the whole body [7]. Around 90% of cancer-related deaths are caused by metastasis of the initial tumor cells to locations away from the central or primary tumor [8]. The proteins of basal lamina are degraded by a mixture of digestive enzymes which secrets from cancer cells and allow it to crawl through. Matrix metalloproteases (MMP) are secreted by cancer cells which cut through the proteins that prevent migrating cancer cells from moving. Tumors can spread to distant organs be through the circulatory system, lymphatic system, and the body wall through the abdominal and chest cavities [9]. A variety of cancer-related genes and molecules have been identified which may lead to cancer progression. Among them the gene family MTA (Metastasis Tumor Antigen) plays a key role in cancer metastasis (Table 1) [10-18]. Gene products produced by MTA genes (i.e. MTA1, MTA2 and MTA3 protein) are closely related to the metastasis cycle [19, 20]. Various factors are regulated by MTA expression including growth factors, growth factor receptors, onco-genes, environmental stress, radiation, inflammation, and hypoxia [21, 22].

**Table 1:**
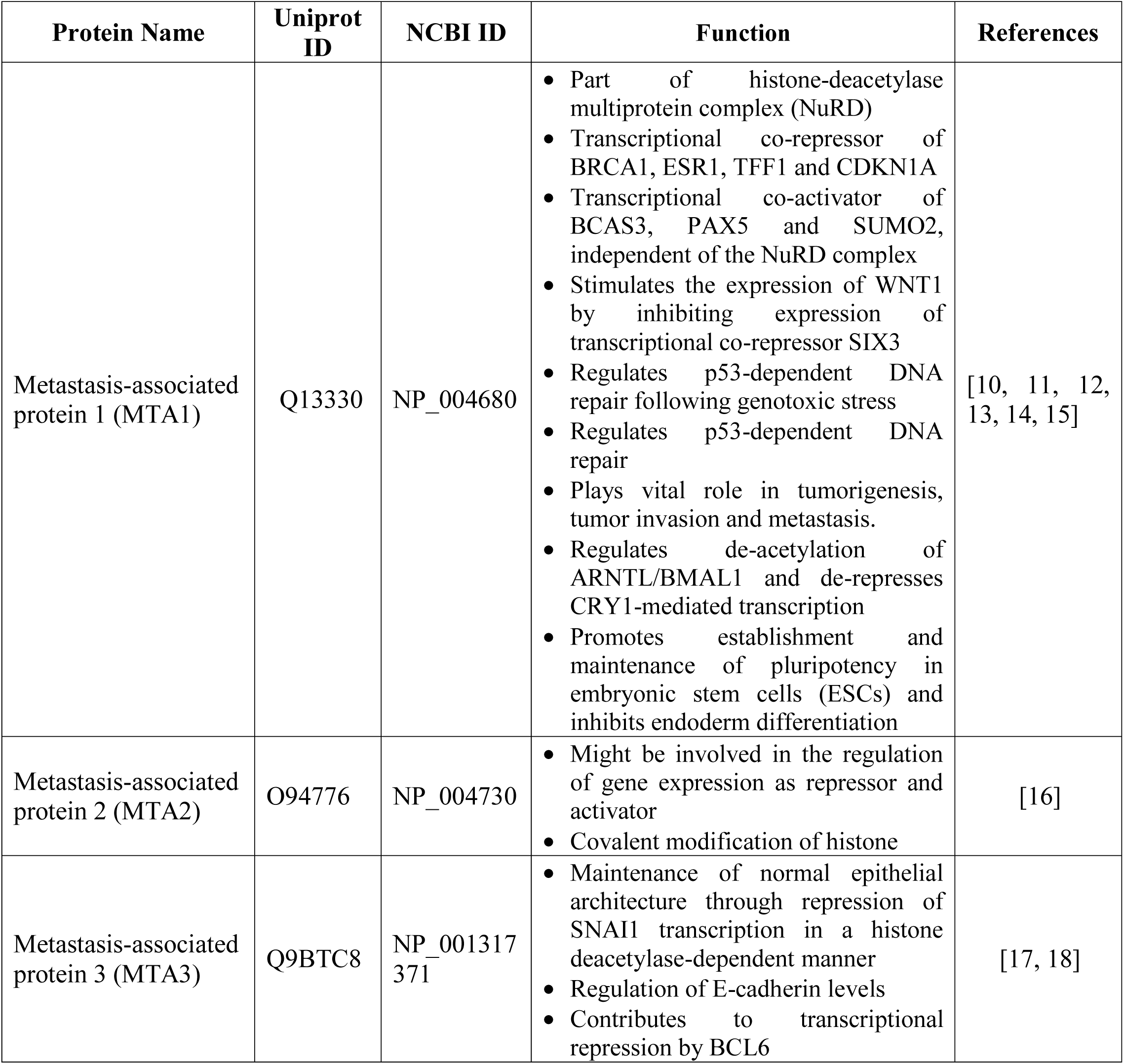
List of metastasis-associated proteins with their identities and functions

The founding member of MTA family, MTA1 is predominantly located in the nucleus and distributed in the extra-nuclear compartments as well [23]. MTA1 and its downstream effectors regulate genes and/or pathways in cancer cells with roles in breast cancer development, invasion, survival, angiogenesis, epithelial-to-mesenchymal transfer, metastasis, DNA damage response and hormone-independence [24]. One of the key cancer promoting activities of MTA1 is its strong interaction with oncogenes in human cancer [25]. It stimulates the transcription of Stat3, breast cancer-amplified sequence 3 [26]. Thus MTA1 helps in cancer progression by inhibiting tumor suppressor genes and stimulating transcription of oncogenes. MTA2 protein is located inside the nucleus and acts in the mechanism chromatin remodeling to control the gene expression. It is over-expressed in human cancer cells and its level of dysregulation is well correlated with more invasive and aggressive phenotypes [27]. MTA2 deacetylates the estrogen receptor alpha and p53 and inhibits their trans-activation functions [28] and thus represses the genes of tumor suppressors and helps in cancer cell development. On the contrary, MTA3 protein locates both in the nucleus and other cellular compartments [29]. The protein controls the selection of targets for nucleosome remodeling and histone deacetylation, thus acting as a transcription repressor. Moreover, MTA family proteins play a key role in aging and Alzheimer’s disease by controlling neuronal genes [30]. Chemical based therapies show various limitations such as drug-resistance, severe side effects, adverse toxicity profile for clinical applications [31, 32]. Natural products, on the other hand, have the potential to form the basis of holistic health care [33]. Due to the medicinal value various plant metabolites are used to treat life-threatening diseases (e.g. cancer, Alzheimer, diabetes, cardiac disease) (Table 2) and to minimize drug toxicity [34-74]. Computing methodologies have been an integral feature of many drug discovery projects, from hits to optimization and beyond. The de-novo detection of active compounds, without choice, can be achieved by virtual screening on protein models, separate from molecular similarity and ligand-based virtual screening [75]. Recent advances in the detection of new ligands and their receptor-bound structures as well as increased results for ligand discovery have rekindled interest in virtual screening that is still generally used for medicinal research [76]. Hence, the study was designed to evaluate some plant-based bioactive compounds and their synthetic analogs to reveal novel therapeutic options against cancer progression by utilizing virtual screening methods and various computational analyses.

**Table 2:**
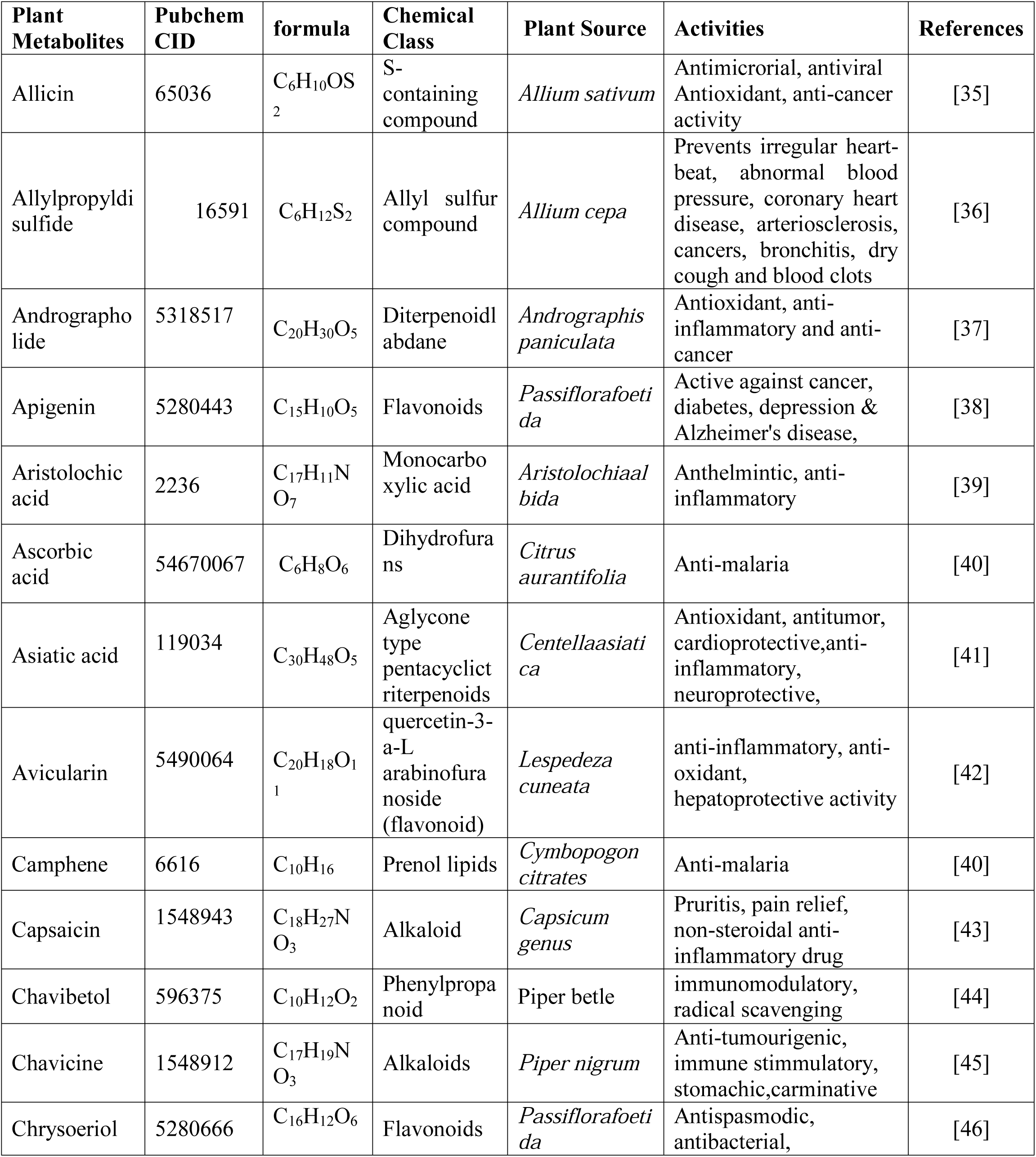
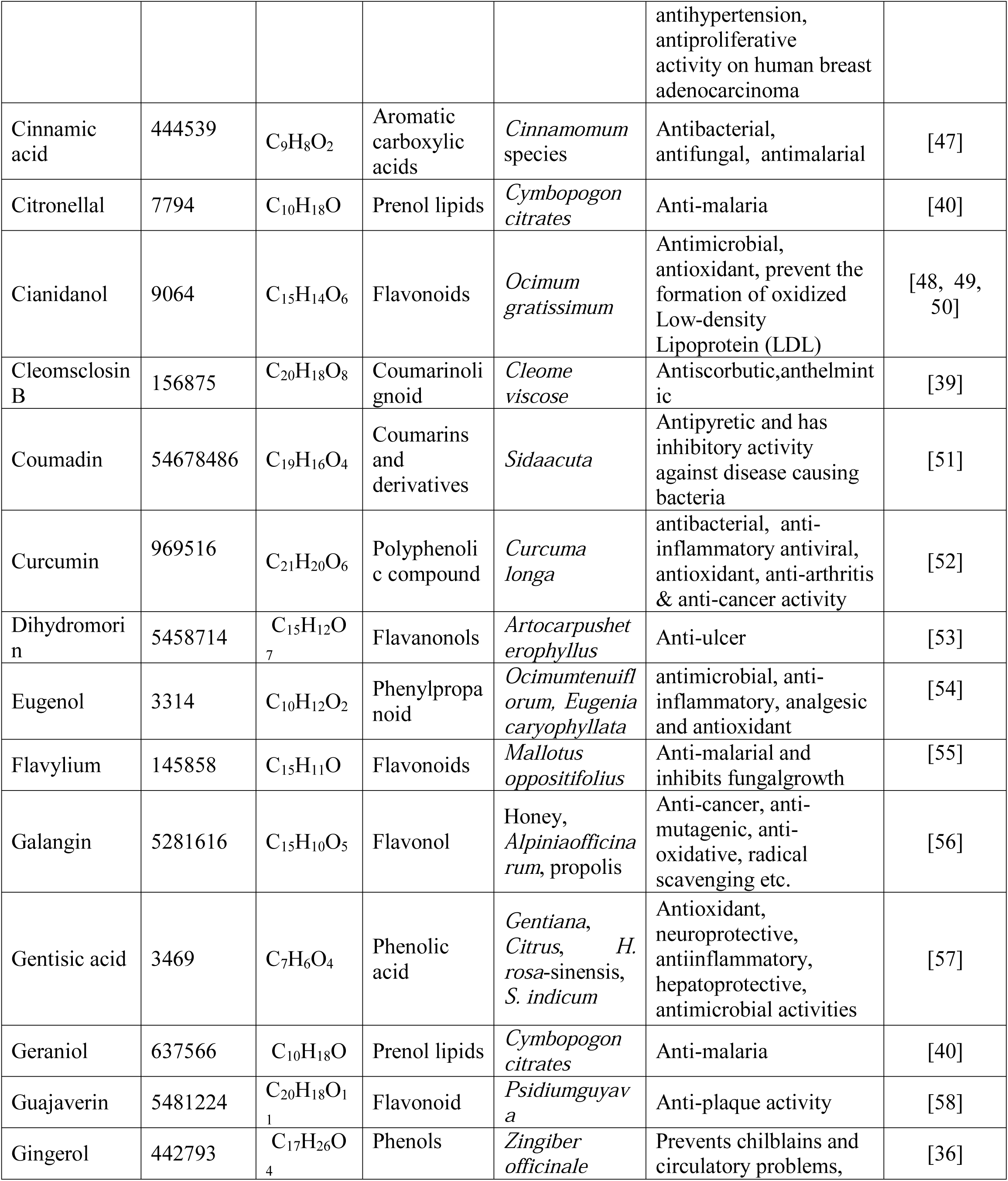
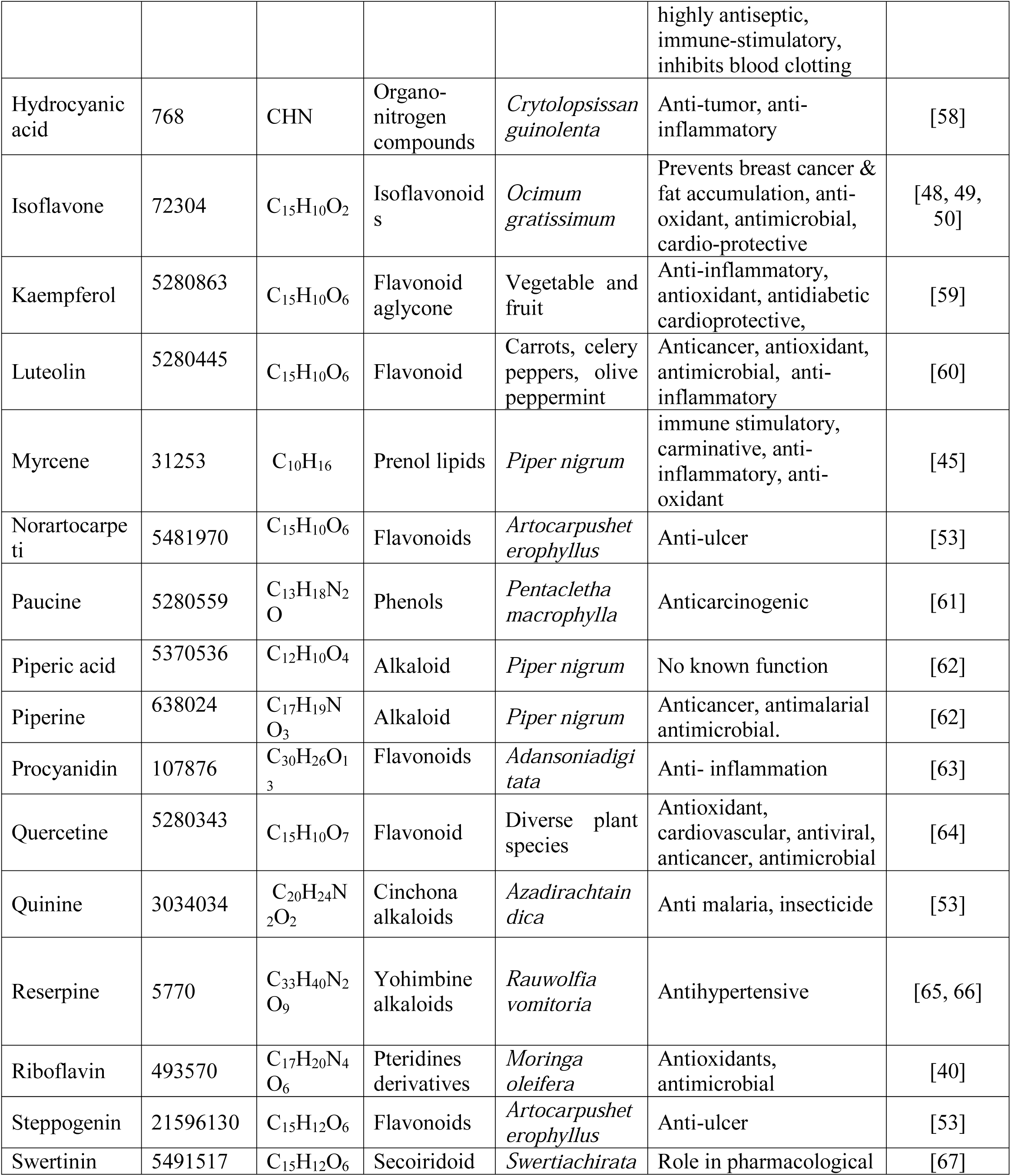
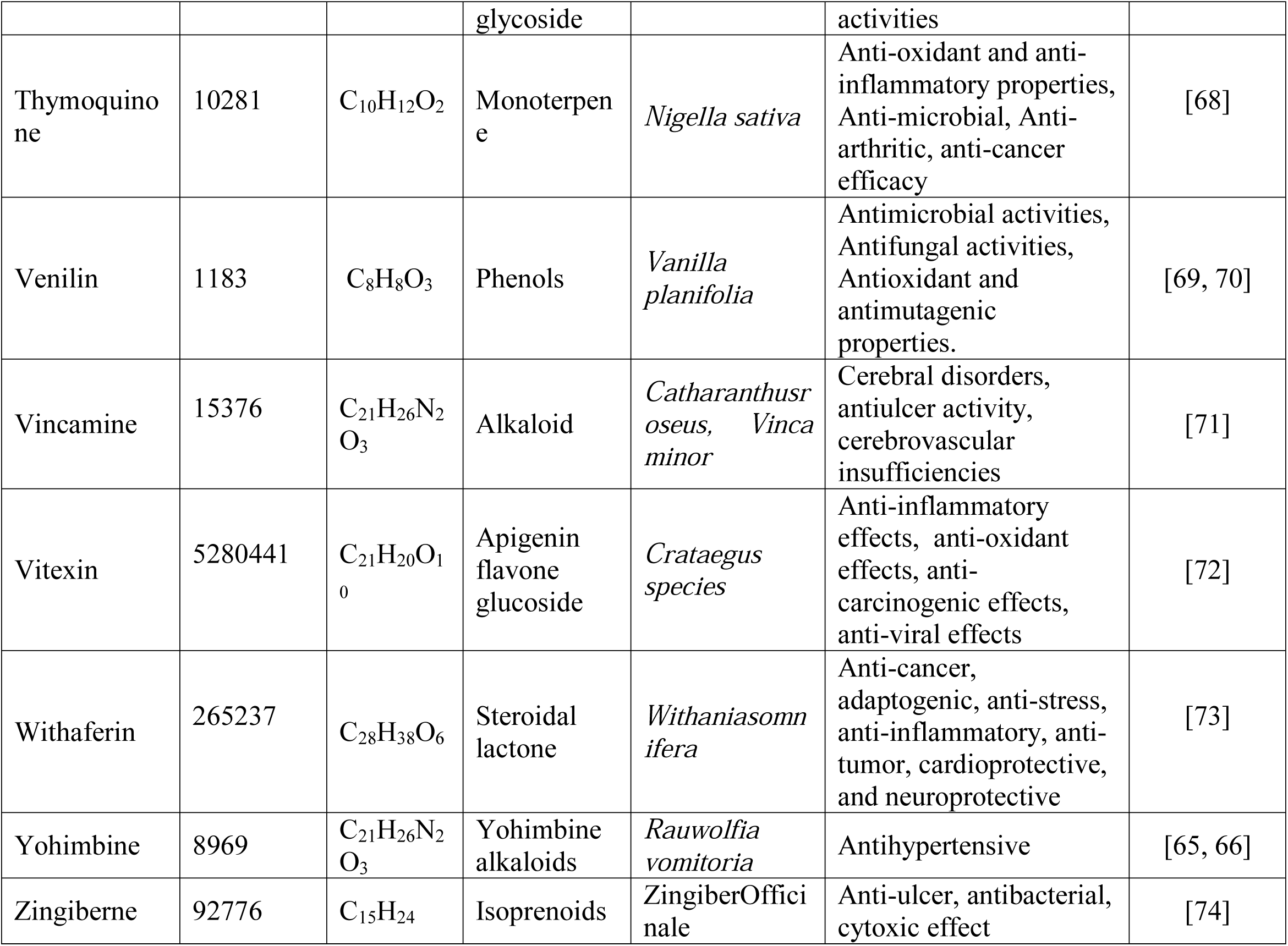
Class, identity, source and function of plant metabolites used in this study

## 2. Materials and methods

### 2.1. Retrieval of metastasis-associated proteins and plant metabolites

The UniProtKB protein database (www.uniprot.org/) was searched for the retrieval of the human metastasis-associated proteins MTA1 (Metastasis Tumor Antigen 1), MTA2 (Metastasis Tumor Antigen 2) and MTA3 (Metastasis Tumor Antigen 3). The proteins were further cross-checked in NCBI (http://www.ncbi.nlm.nih.gov/) protein database for the corresponding accession numbers. Besides, a total of 50 plant metabolites belonging to different classes were retrieved from PubChem server in SDS (3D) format [77]. OpenBabel v2.3 software was used to convert the structures into PDB format [78, 79].

### 2.2. Analysis of physicochemical properties and sub-cellular localization

Various physicochemical properties of the MTA proteins were demonstrated using ProtParam, a tool from ExPASy server [80]. Molecular weight, theoretical pI, extinction co-efficient, grand average of hydropathy (GRAVY), half-life, aliphatic index (AI), instability index and amino acid composition were calculated. For understanding protein function, it is important to find out the sub-cellular localization of proteins. CELLO [81], ngLOC [82] and PSIpred [83] servers were used for this purpose.

### 2.3. Secondary structure prediction and functional analysis

Secondary structures, the conformation that is adopted by the polypeptide backbone of a protein, are composed of regions stabilized by hydrogen bonds between different atoms [84]. Different secondary structural elements (e.g. helices, pleated sheets, turns) were analyzed using CFSSP (Chou and Fasman secondary structure prediction) server [85] and PSIPRED (PSI-blast based secondary structure prediction) tools [86]. Families and super-families of the MTA proteins were analyzed using SUPERFAMILY v2.0 [87, 88]. Moreover, NCBI-CDD (NCBI Conserved Domains Database), a protein annotation resource of NCBI [89] and MOTIF Search server [90] determined the conserved domains and motifs of these proteins, respectively.

### 2.4. Tertiary structure prediction, refinement and validation

All three proteins (i.e. MTA1, MTA2, MTA3) were subjected to 3D modeling via I-TASSER (Iterative Threading Assembly Refinement) through using a hierarchical protein structure prediction approach [91]. The unwanted errors in the predicted models were removed by GalaxyRefine service of GalaxyWEB server [92]. The best refined models of the studied proteins were determined through quality assessment and validation. Commonly used tools, ERRAT [93] and RAMPAGE [94] were employed to meet this purpose. The SAVES v5.0 (https://servicesn.mbi.ucla.edu/SAVES/) server was used to observe amino acid distributions within studied protein models.

### 2.5. Active site prediction and mobility analysis

Active sites identification is a crucial step in drug discovery which evaluates the size of an active site, the number and properties of sub-sites and details of binding interaction [95]. CASTp (computed atlas of surface topography of proteins) server predicted the active sites in the refined models of three MTA proteins [96]. iMODs server represents the collective motion of proteins by evaluating the normal modes (NMA) in internal coordinates [97]. It predicted the extent as well as direction of the inherent motions of studied proteins. The deformability, eigen values and covariance-matrix were also analyzed.

### 2.6. Screening of plant metabolites against MTA proteins and analysis of drug surface hotspots

Molecular Docking is a beneficial tool to perform virtual screening on various compounds and to infer how the ligands inhibit their targets [98]. The binding affinities of 50 plant metabolites with 3 MTA proteins were determined using PatchDock server [99]. In this study, Metarrestin (Pubchem CID: 50985821) was used as positive controls for their inhibitory features against MTA proteins. The docked complexes were refined via FireDock refinement tool [100]. Visualization of ligand binding complexes was performed by Discovery Studio v3.1 [101] and PyMOL v2.0 software [102].

### 2.7. Molecular dynamics study

Stability of the docked complex was predicted by molecular dynamics study. The structural dynamics of the studied proteins were assessed using iMOD server for its quick and effective measurements than other molecular dynamics (MD) simulations tools [103; 104]. The structural communications fingerprints in the biomolecules were measured through exploring global metapath and path distribution via WebPSN server [105]. Moreover, the protein dynamics as like the protein contact maps for simulation trajectory, fluctuation plots were analyzedby CABS-flex 2.0 server, CABS-flex is more suited to detectnon-obvious dynamic fluctuations within, for example, the biologically important, well-defined secondary structural elements [106].

### 2.8. Drug profile analysis of top metabolites

Absorption, distribution, metabolism, and excretion (ADME) properties are significant pharmacological features to facilitate drug development processes for safe and effective therapeutics production [107]. SwissADME server was used to evaluate ADME properties of top plant metabolites [108]. To assess the Blood-brain barrier (BBB) in the studied compounds, BOILED-Egg model was applied [109]. Moreover, pkCSM, an online tool was prioritizing to evaluate the relative toxicity of top metabolites [110, 111].

### 2.9. Prediction of drug targets and available drug molecules from drugbank

SwissTargetPrediction tool was used to assume the macromolecular drug targets of top metabolites [112]. Furthermore, SwissSimilarity web tool was employed to identify already approved drug molecules with similar structure from DrugBank [113]. The server performed ligand-based virtual screening of several libraries of small molecules to trace approved, experimental and commercially available drugs from DrugBank [113].

## 3. Results

### 3.1. Retrieval of metastasis-associated proteins and plant metabolites

UniProt entry Q13330 (715 amino acids), O94776 (668 amino acids) and Q9BTC8 (594 amino acids) corresponding to MTA1, MTA2 and MTA3 respectively, were retrieved and cross-checked with NCBI protein database accession numbers (i.e. NP_004680.2, NP_004730.2 and NP_001317371.1 respectively) (Table 1). PubChem ID, molecular weight, formula, source, function and others properties of retrieved 50 metabolites were listed in Table 2.

### 3.2. Analysis of physicochemical properties and sub-cellular localization

The molecular weight of the metastasis-associated proteins ranged from 67.5to 80.8 kDa. The isoelectric points were predicted between 8.80 and 9.70, suggesting that the proteins are basic in nature. Moreover, aliphatic index around 75 revealed the proteins as thermo-stable. The negative GRAVY values indicated that the proteins will have a good interaction with water (Table 3). The localization of the MTA proteins (i.e. MTA1, MTA2, MTA3) were predicted as nuclear protein.

**Table 3:** Physicochemical property analysis of MTA proteins

| Proteins | Formula | Molecular weight | p<sup>I</sup> | Extinction coefficient | Half life | Aliphatic index | GRAVY |
| --- | --- | --- | --- | --- | --- | --- | --- |
| MTA 1 | C<sub>3565</sub>H<sub>5658</sub>N<sub>1032</sub><br>O<sub>1055</sub>S<sub>29</sub> | 80786.16 | 9.34 | 0.1% | 30 hours<br>(mammalian<br>reticulocytes, *in vitro*)<br>>10 hours (*E. coli, in vivo*) | 75.48 | -0.617 |
| MTA 2 | C<sub>3303</sub>H<sub>5303</sub>N<sub>969</sub><br>O<sub>969</sub>S<sub>29</sub> | 75023.09 | 9.70 | 0.1% | 30 hours<br>(mammalian<br>reticulocytes, *in vitro*).<br>>10 hours (*E. coli, in vivo*). | 77.19 | -0.528 |
| MTA 3 | C<sub>2982</sub>H<sub>4693</sub>N<sub>837</sub><br>O<sub>900</sub>S<sub>26</sub> | 67503.69 | 8.80 | 0.1% | 30 hours<br>(mammalian<br>reticulocytes, *in vitro*)<br>>10 hours (*E. coli, in vivo*). | 74.75 | -0.541 |

### 3.3. Secondary structure prediction and functional analysis

Results showed that MTA1 had 62.9% (450 residues) helix, 30.8% (220 residues) sheet and 15% (107 residues) turn, while MTA2 showed to have 68.9% (460 residues) helix, 37.0% (247 residues) sheet and 13.6% (91 residues) turn regions. Similarly, MTA3 exhibited 63.3% (376 residues) helix, 44.4% (264 residues) sheet and 15.5% (92 residues) turns. Higher numbers of α-helices are suggesting these proteins as thermostable (Figure 1) [114]. All of the studied proteins comprised one Homeodomain-like superfamily and one Myb/SANT domain family (Figure 2). Protein annotation resource NCBI-CDD revealed that these MTA proteins have five domains, namely Bromo Adjacent Homology domain (BAH MTA), MTA R1 domain, Myb-Like DNA-Binding Domain (SANT MTA3 like domain), zinc finger binding to DNA consensus sequence [AT]GATA[AG] (ZnF_GATA) and the ELM2 (Egl-27 and MTA1 homology 2) domain (Figure 2). Only MTA2 had an additional domain named PHA03247 super family that is large tegument protein UL36.MotifFinder predicted six motifs, MTA_R1, BAH, ELM2, GATA, Myb_DNA-binding and Myb_DNA-bind_7 for each protein. Again, MTA2 showed an extra motif named Cytochrome C_3_ (Figure 2B).

**Figure 1:**
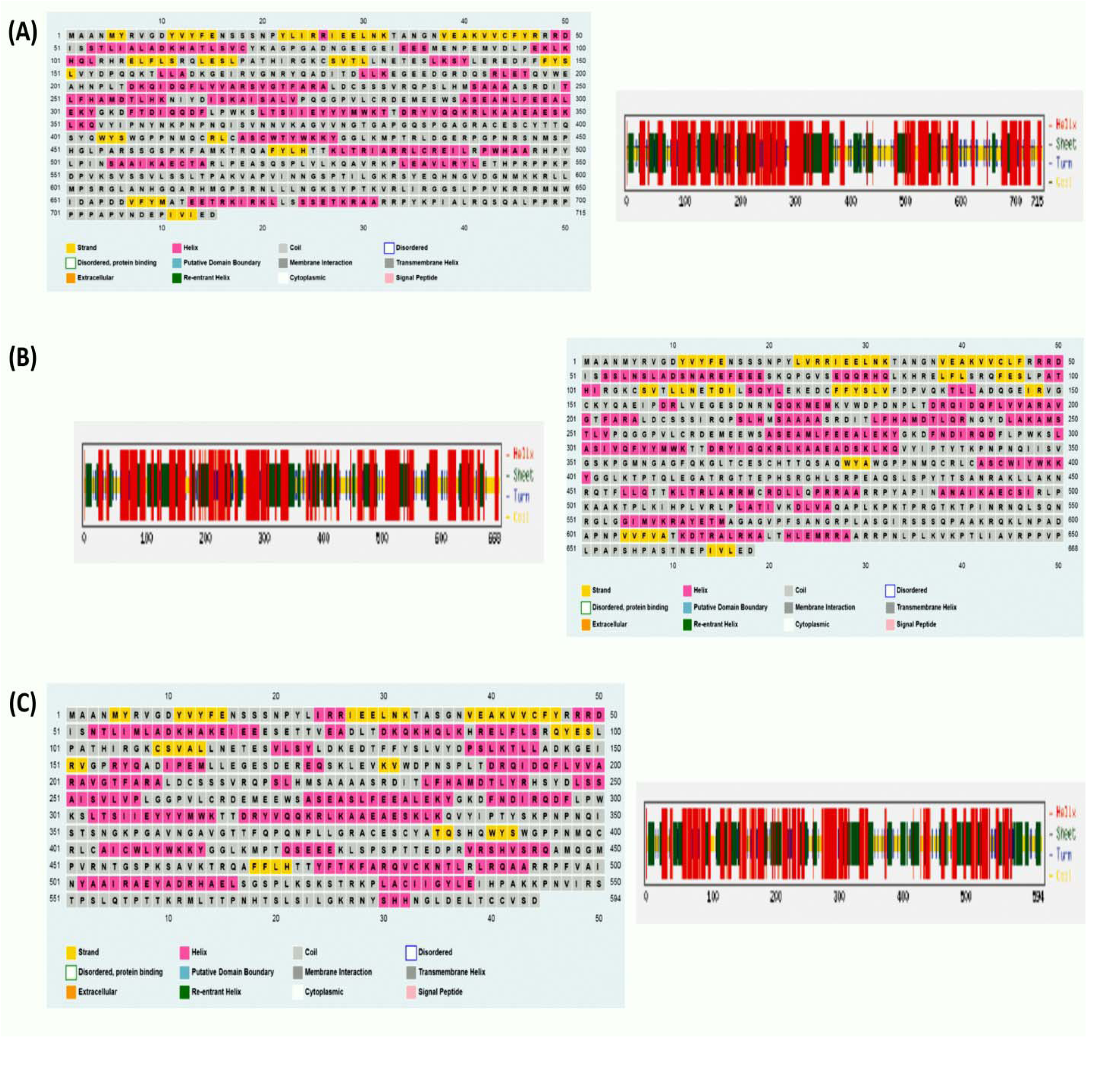
Predicted secondary structure of MTA1 (A), MTA2 (B) and MTA3 protein (C).

**Figure 2:**
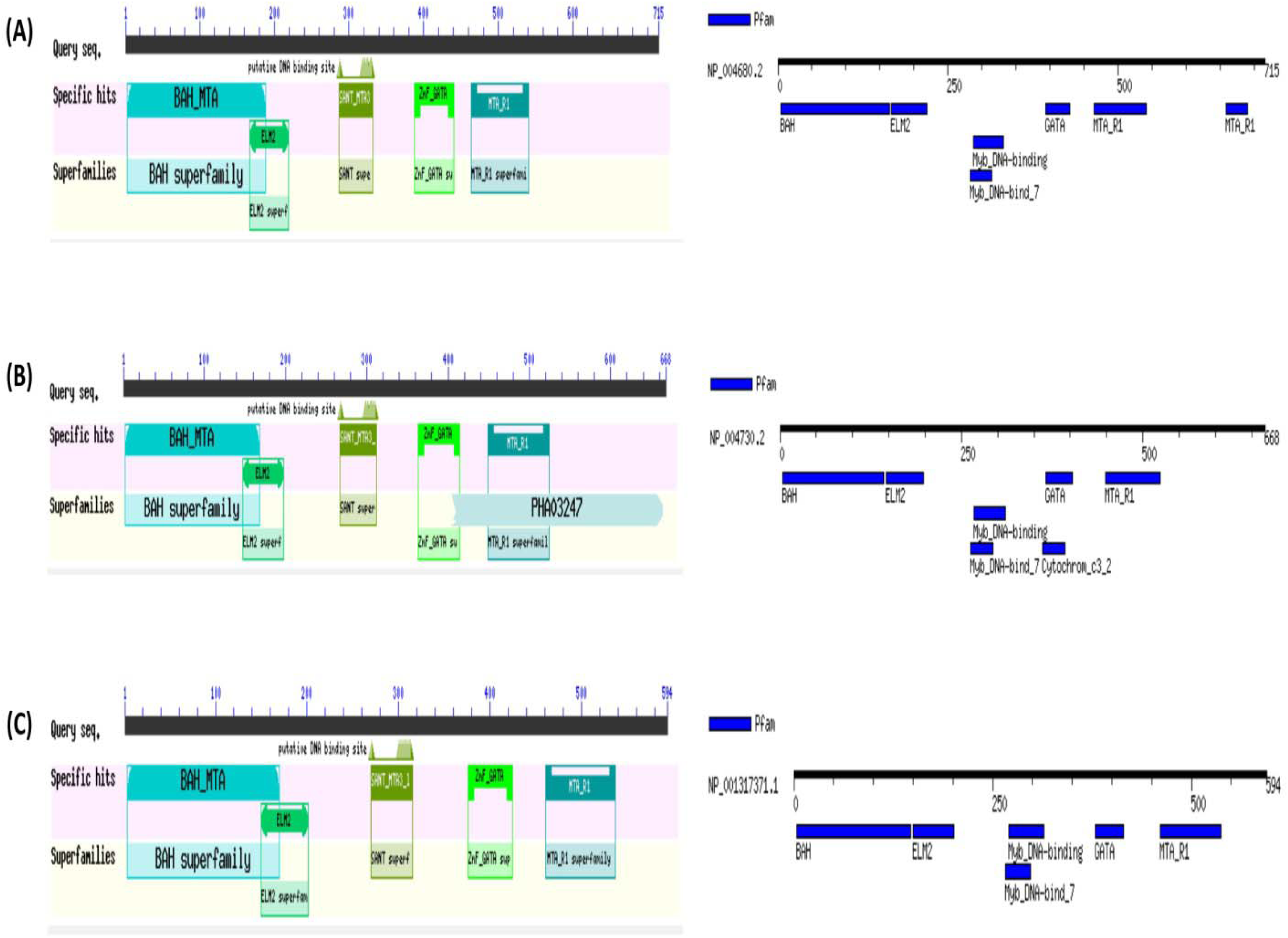
Predicted domains and motifs of MTA1 (A), MTA2 (B), MTA3 (C).

### 3.4. Tertiary structure prediction, refinement and validation

I-TASSER server performed three dimensional homology modeling and predicted five models for each protein. The best models were identified based on C-Score (Figure 3-5A), which showed overall quality factors 79.914(MTA1), 80.793(MTA2) and 85.665 (MTA3) at 0.01 and 0.05 level of significance (Figure 3-5B). The results of Ramachandran plot analysis revealed 87.2% residues in favored, 9% residues in allowed and 3.8%residues in the outlier region for MTA1. The refined models of MTA2 and MTA3 showed 88.4% and 91.2% residues in the favored region, while only 1.8% and 1.9% residues were in the outlier region, respectively (Figure 3-5C). Besides, the amino acid distributions of the refined models were revealed by comparing expected and observed structure through SAVES server (Figure 3-5D).

**Figure 3:**
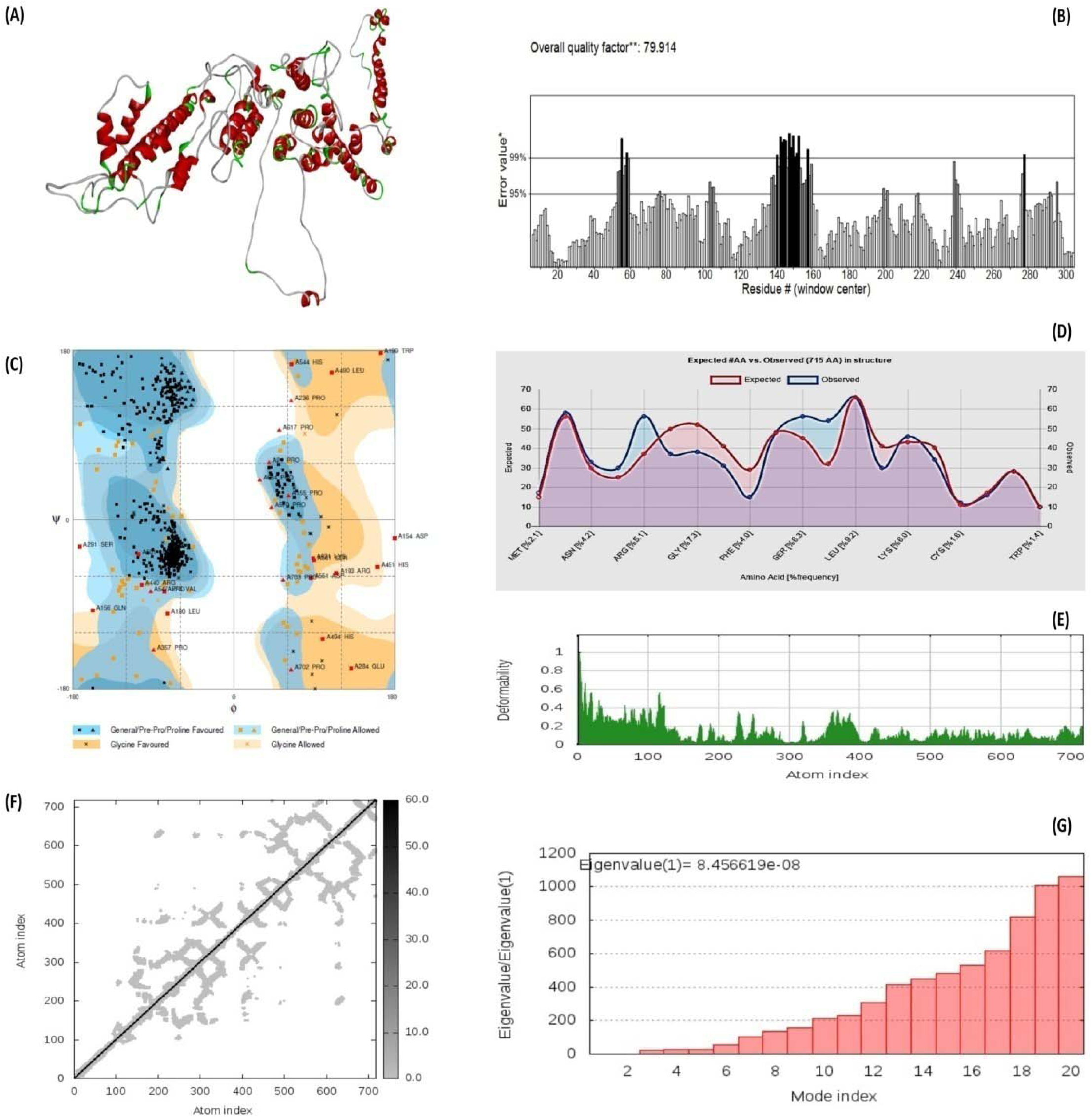
Homology modeling (A), quality factor (B), ramachandra plot assessment (C), amino acid Distribution plot (D), deformability (E), atom index (F), eigen value (G) of MTA1 protein.

**Figure 4:**
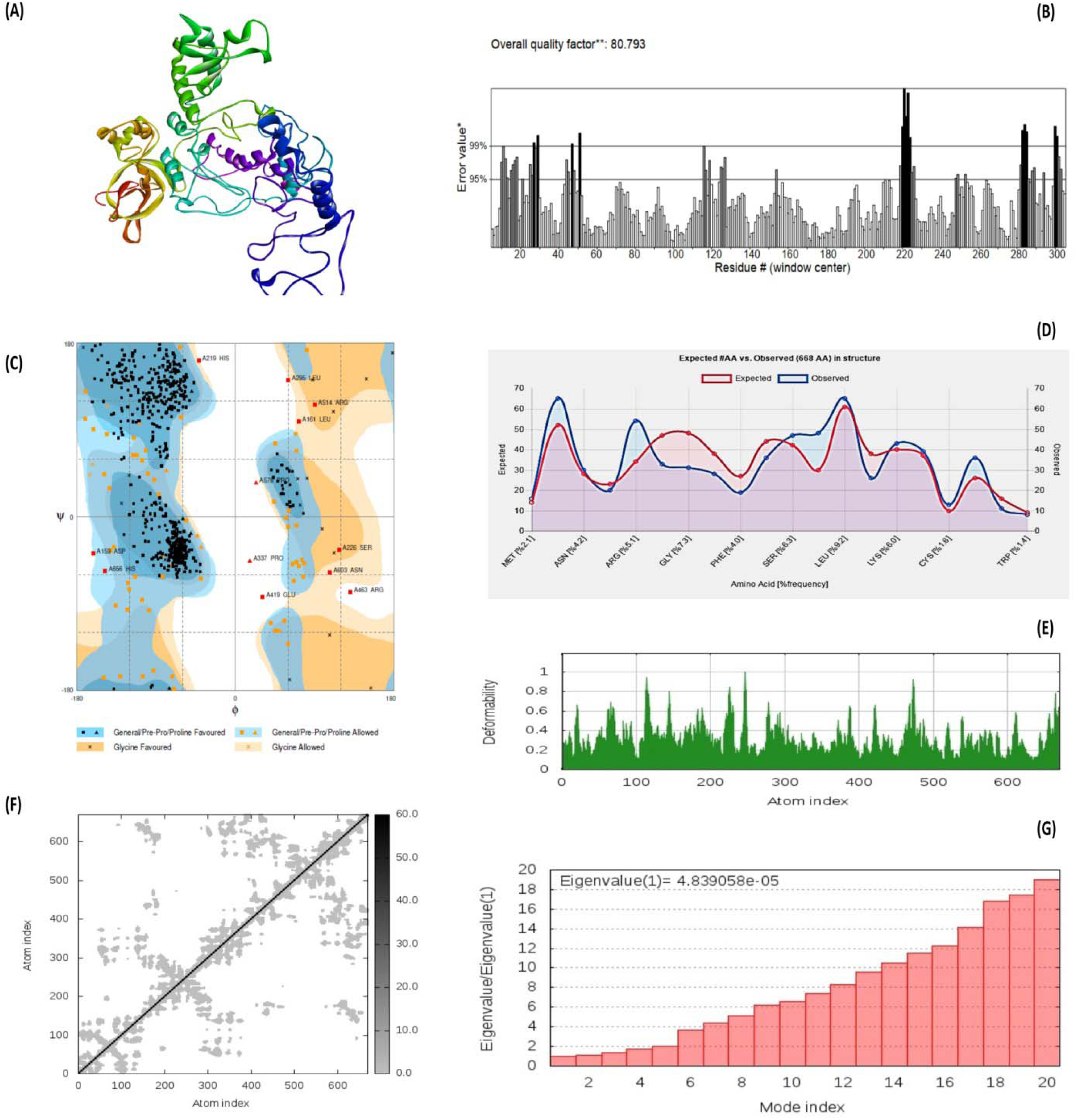
Homology modeling (A), quality factor (B), ramachandra plot assessment (C), amino acid Distribution plot (D), deformability (E), atom index (F), eigen value (G) of MTA2 protein.

**Figure 5:**
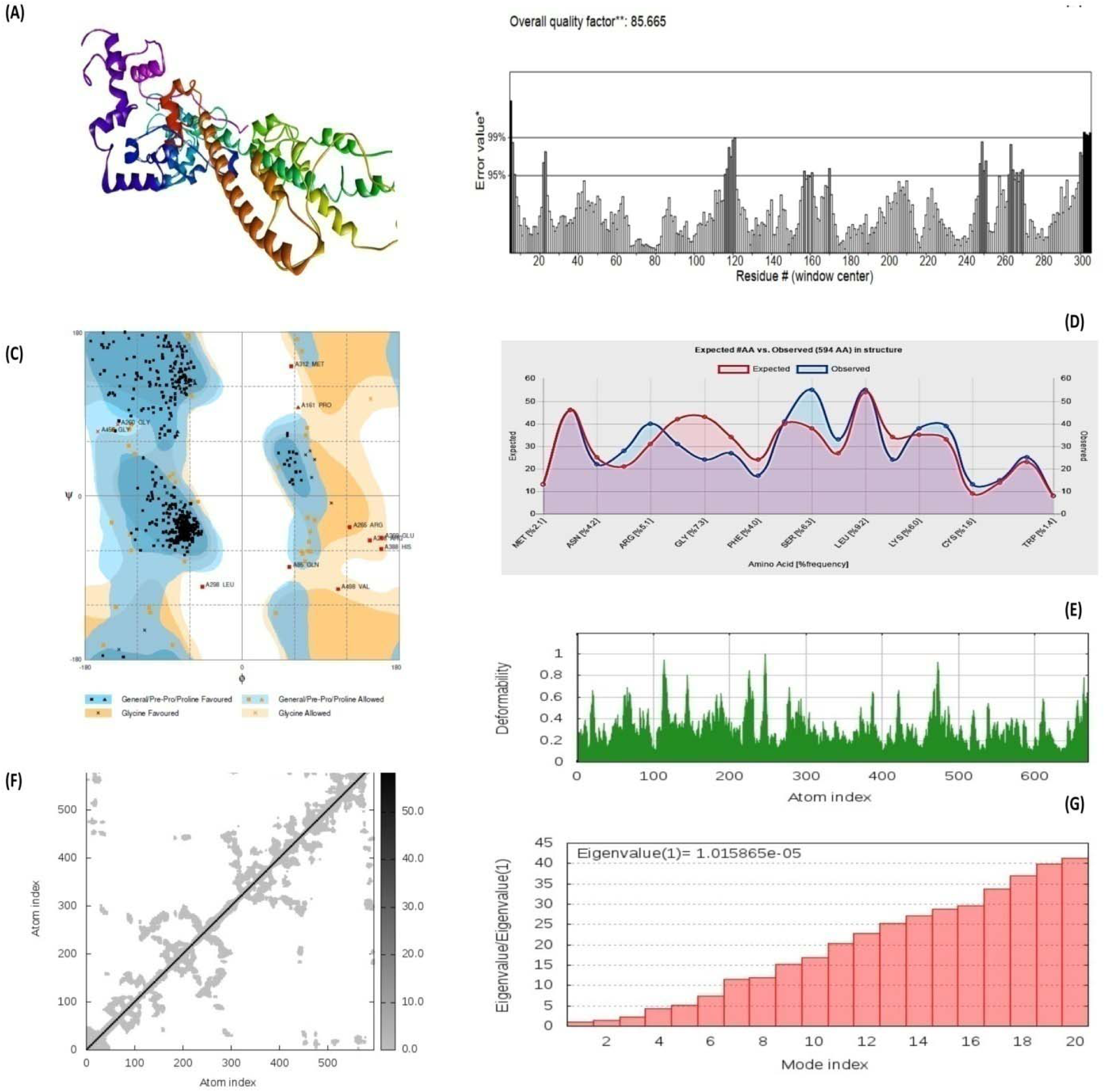
Homology modeling (A), quality factor (B), ramachandra plot assessment (C), amino acid Distribution plot (D), deformability (E), atom index (F), eigen value (G) of MTA3 protein.

### 3.5. Active site prediction and mobility analysis

CASTp server revealed 135, 138 and 93 active sites for refined models of MTA1, MTA2 and MTA3 (Supplementary file 1). The best pockets showed an area and volume of 1318.734 (SA) and 557.492 (SA) for MTA1; 901.007 (SA) and 1563.224(SA) for MTA2; 4094.143 (SA) and 3963.371 (SA) for MTA3 protein (Figure 6). Stability of the modeled structures was determined in terms of deformibility, eigenvalue and covariance matrix. The main-chain deformability of the MTA proteins was negligible as indicated by hinges in the chain (Figure 3-5E). The higher eigenvalues of MTA1 (8.456619e^-08^), MTA2 (8.839058e^-05^) and MTA3 (1.015865e^-05^) are representative of higher energy which is required to deform the protein structures (Figure 3-5G). The elastic network models as shown in Figure 3-5F; defined the pairs of atoms connected by springs, where dots are colored according to the degree of stiffness.

**Figure 6:**
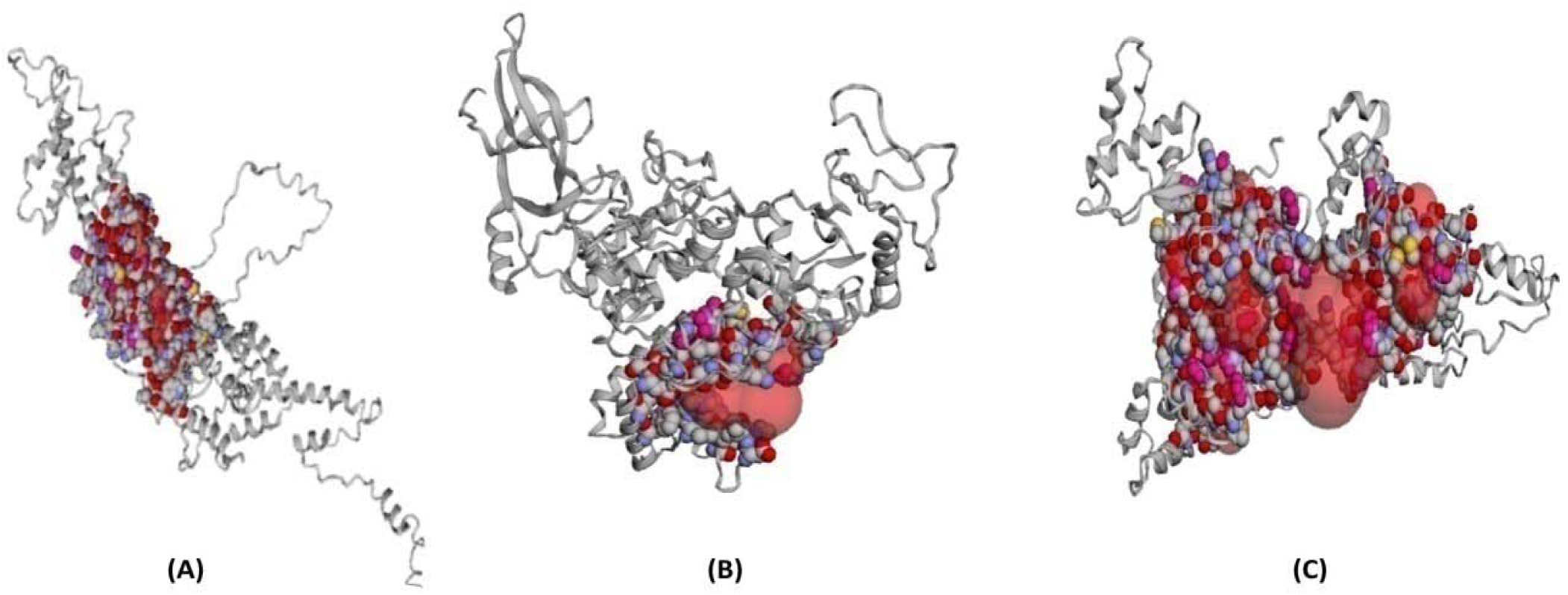
Predicted ligand binding pockets of MTA1 (A), MTA2 (B), MTA3(C) via CASTp.

### 3.6. Screening of plant metabolites against MTA proteins and analysis of drug surface hotspots

The MTA proteins (macromolecules) and plant metabolites (ligands) were used for molecular docking to assess the affinity between the ligands and macromolecules. Top metabolites were ranked according to the global binding energy (Supplementary file 2). Notably, isoflavone, gingerol, citronellal and asiatic acid showed higher binding affinity with each of the three proteins and featured in the top list (Table 5). Isoflavone showed maximum binding interaction with MTA2 (−54.70 kcal/mol) and MTA3 (−69.20kcal/mol) (Figure 7B and 7C; Table 5). Gingerol also experienced minimum binding energy with MTA1 (−51.23kcal/mol), MTA2 (−52.71 kcal/mol) and MTA3 (−48.02 kcal/mol) protein (Figure 7A, Table 5).The ligand binding interactions and structural conformations of each protein were investigated to unravel the drug surface hotspots. Results revealed that the regions from 433 to 498 and 629 to 633 were crucial binding sites for MTA1 protein, where Pro629, Lys631, Val632, Arg633 positions were most dominant. Again, residues of 333-397 and 616-624 regions were identified as top surface hotspot for MTA 2. Moreover, two residues i.e. Lys618 and Ala619 were involved to form the docked complexes in maximum cases. The ligands showed maximum binding affinity for the region of 425 to 458 amino acid sequences in case of MTA3 protein.

**Table 4:** Functional domains and motif analysis of MTA proteins

| Protein | Pfam | Position (E-value) | Description |
| --- | --- | --- | --- |
| MTA1 | MTA_R1 | 464..541 (3.9e-34)<br>658..690 (0.52) | PF17226, MTA R1 domain |
|  | BAH | 4..164 (1.3e-35) | PF01426, BAH domain |
|  | ELM2 | 167..219 (1.7e-12) | PF01448, ELM2 domain |
|  | GATA | 393..429 (2e-09) | PF00320, GATA zinc finger |
|  | Myb_DNA-binding | 287..33 (6.5e-06) | PF00249, Myb-like DNA-binding domain |
|  | Myb_DNA-bind_7 | 283..314 (0.0079) | PF15963, Myb DNA-binding like |
| MTA2 | MTA_R1 | 448..524 (4.4e-28) | PF17226, MTA R1 domain |
|  | BAH | 4..143 (2.9e-28) | PF01426, BAH domain |
|  | ELM2 | 147..198(4.9e-15) | PF01448, ELM2 domain |
|  | GATA | 367..403 (2.6e-11) | PF00320, GATA zinc finger |
|  | Myb_DNA-binding | 267..311 (0.00011) | PF00249, Myb-like DNA-binding domain |
|  | Myb_DNA-bind_7 | 263..294 (0.031) | PF15963, Myb DNA-binding like |
|  | Cytochrom_c3_2 | 362..393 (0.39) | PF14537, Cytochrome c3 |
| MTA3 | BAH | 4..147 (1.9e-31) | PF01426, BAH domain |
|  | MTA_R1 | 461..537 (1.7e-28) | PF17226, MTA R1 domain |
|  | ELM2 | 150..201 (1.7e-15) | PF01448, ELM2 domain |
|  | GATA | 379..415 (4e-09) | PF00320, GATA zinc finger |
|  | Myb_DNA-binding | 270..314 (1.8e-05) | PF00249, Myb-like DNA-binding domain |
|  | Myb_DNA-bind_7 | 266..297 (0.016) | PF15963, Myb DNA-binding like |

**Table 5:**
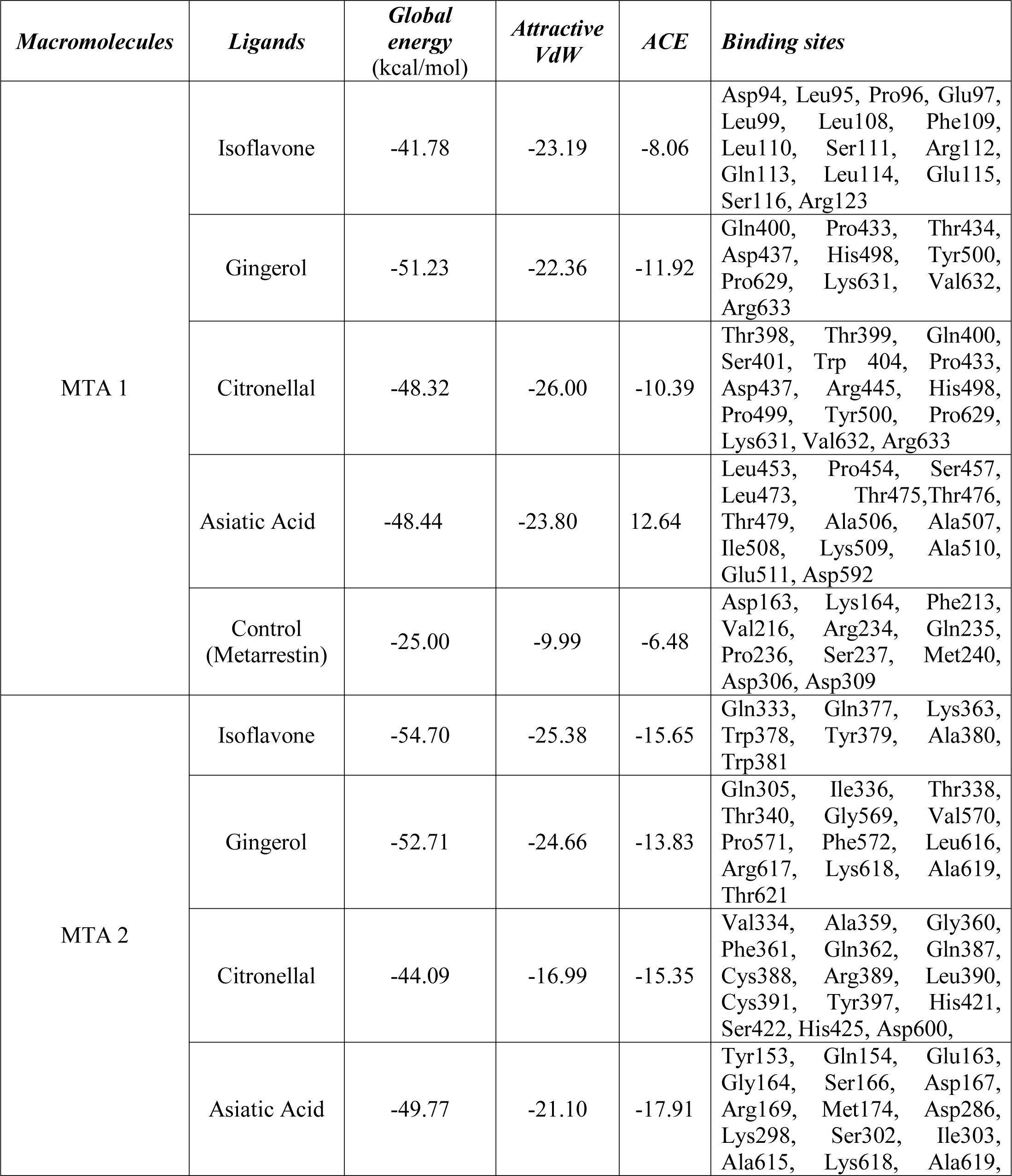
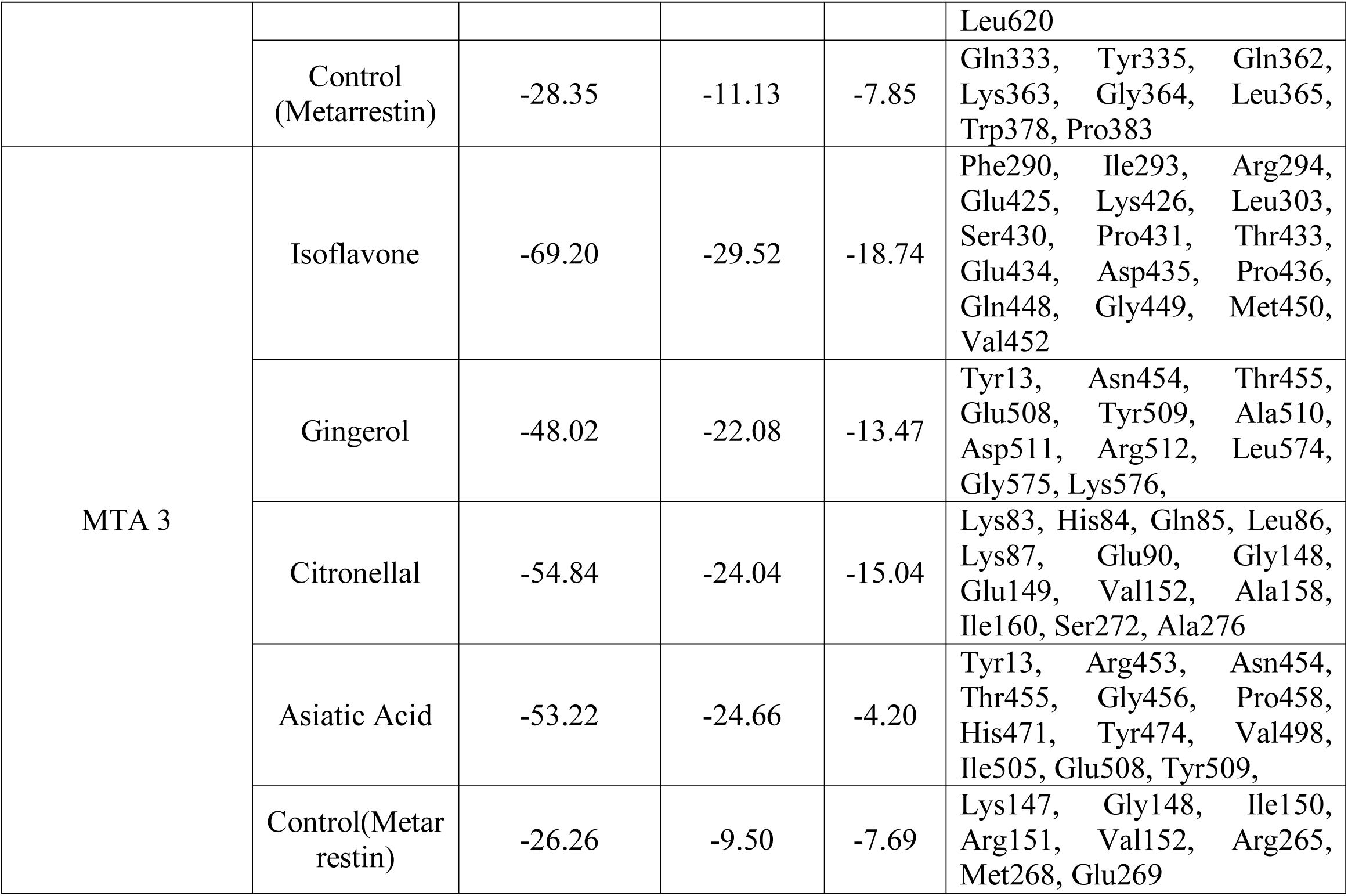
Docking results of top metabolites against MTA proteins

**Figure 7:**
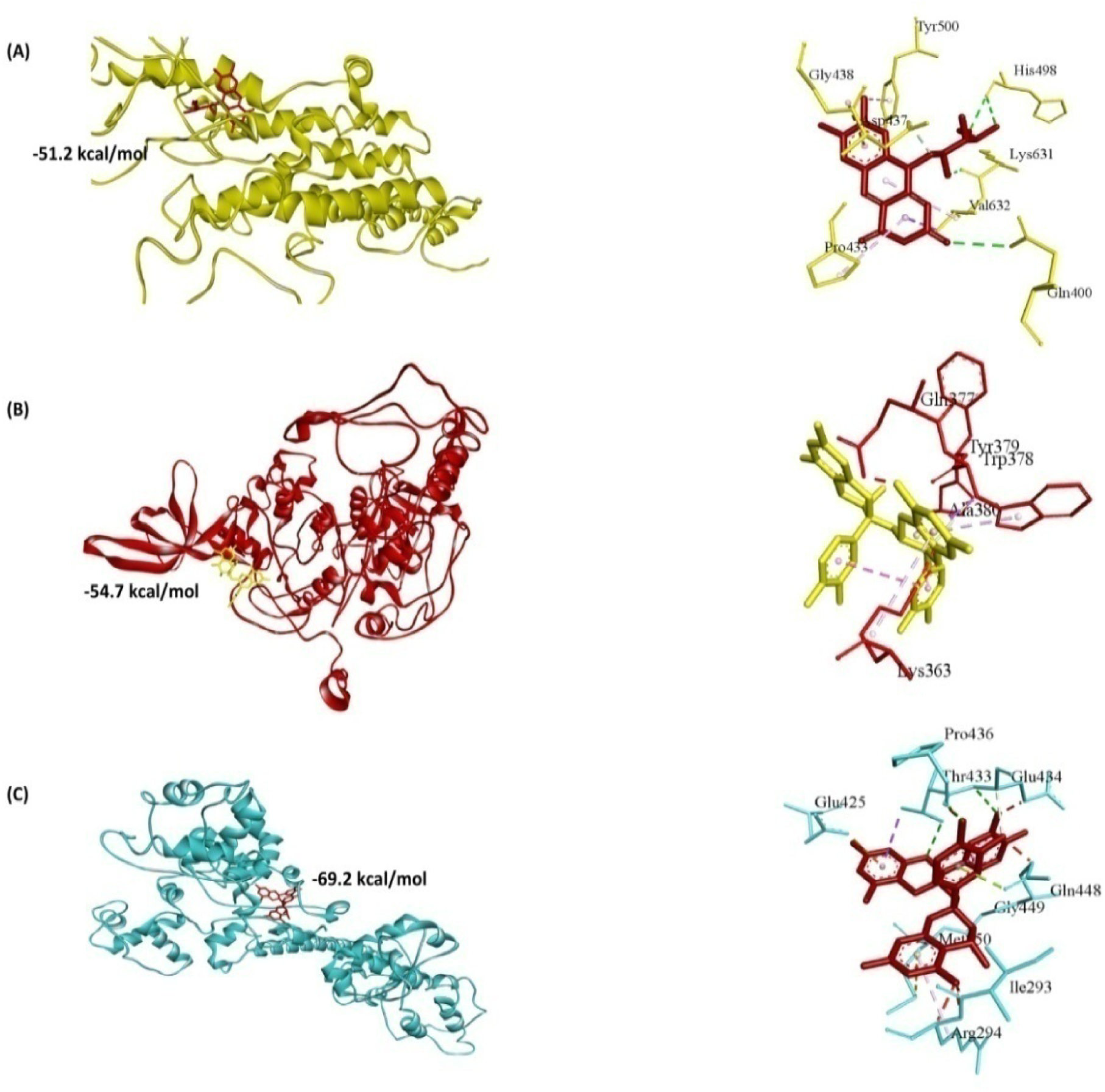
Molecular interaction of gingerol with MTA1 protein (A), isoflavone with MTA 2 (B) and isoflavone with MTA3 protein (C).

### 3.7. Molecular dynamics study

The iMODS server conducted such an analysis taking into account the complex molecule’s internal coordinates, the B-factor values obtained from NMA are equal to RMS (Figure 10).The interaction of MTA1 and Gingerol revealed the total number of nodes involved in the recurrent links was 102. The highest average correlation between each node was 1.57067 with an average path correlation of 1.57067 and the total hubs determined were 90, while the average percentage of hub nodes present in the global pool was 45.09% (Figure 10A). Additionally, the average of the interaction strength among the links present in the global pool was designated by path force 5.82366. The meta path generated from the interactions of isoflavone with MTA2 and MTA3 revealed 116,84 and 115,83 nodes respectively (Figure 10B and Figure 10C). Maximum path correlation was found 1 for both cases with an average path correlation 0.9686689 and 0.974525, respectively for MTA1 and MTA2. The average path force was 6.7 in MTA1 and 7.15 in case of MTA2. The protein dynamics analysis of complexes provided by the CABS-flex 2.0 server insights the flexible and rigid portions of the complexes. The MTA1-gingerol, MTA2-Isoflavone and MTA3-isoflavone complexes had lesser fluctuations in top integrated best 10 models (Figure 11). The root mean square fluctuation plot insights the MTA1-Gingerol, MTA2-isoflavone and MTA3-isoflavone complex’s fluctuation ranged 0-16 Å, 0-7 Å, 0-5.5 Å respectively (Figure 11).

**Figure 8:**
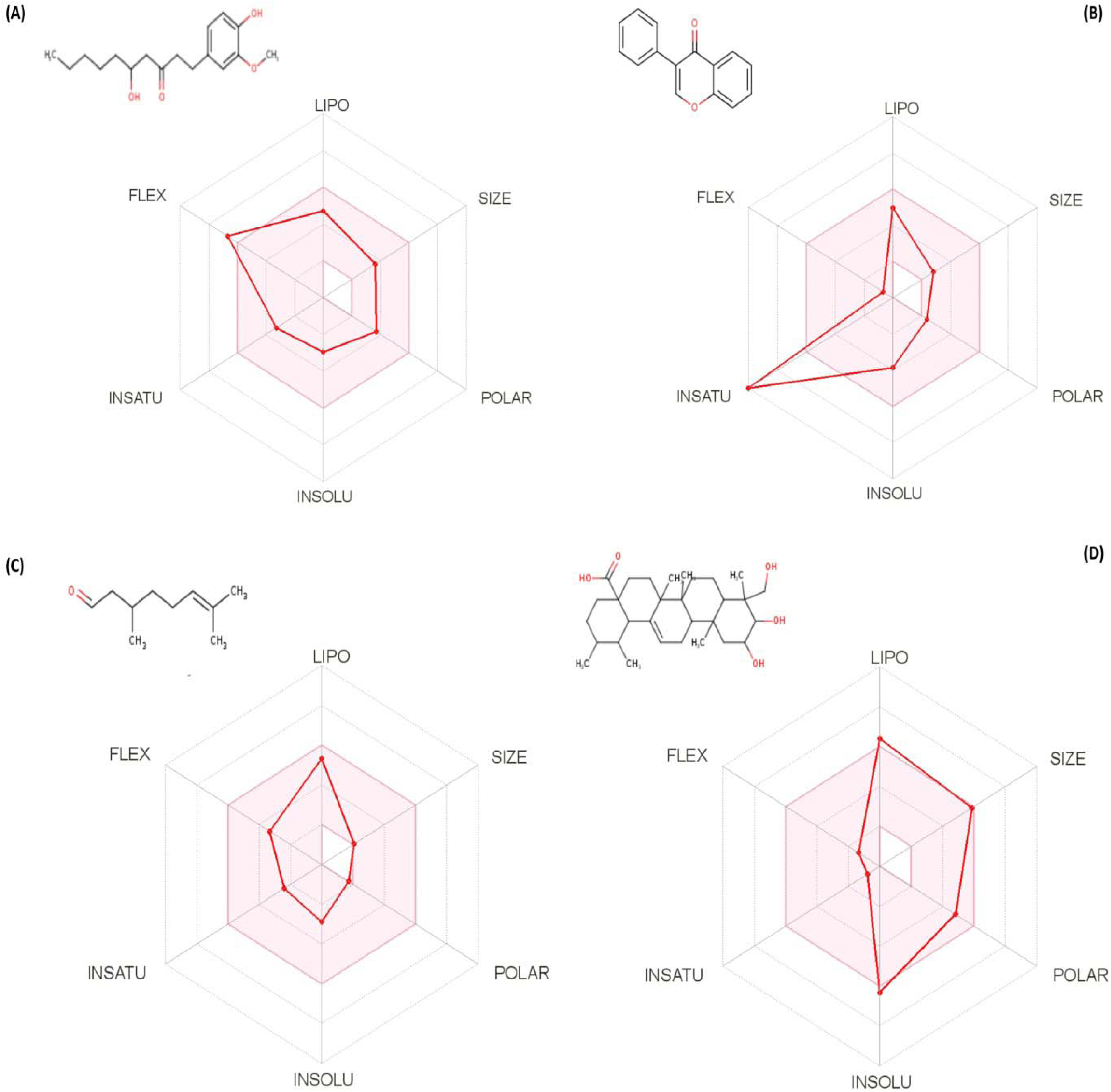
Chemical structure and ADME analysis of top drug candidates; (A) gingerol, (B) isoflavone, (C) citronellal, (D) asiatic acid.

**Figure 9:**
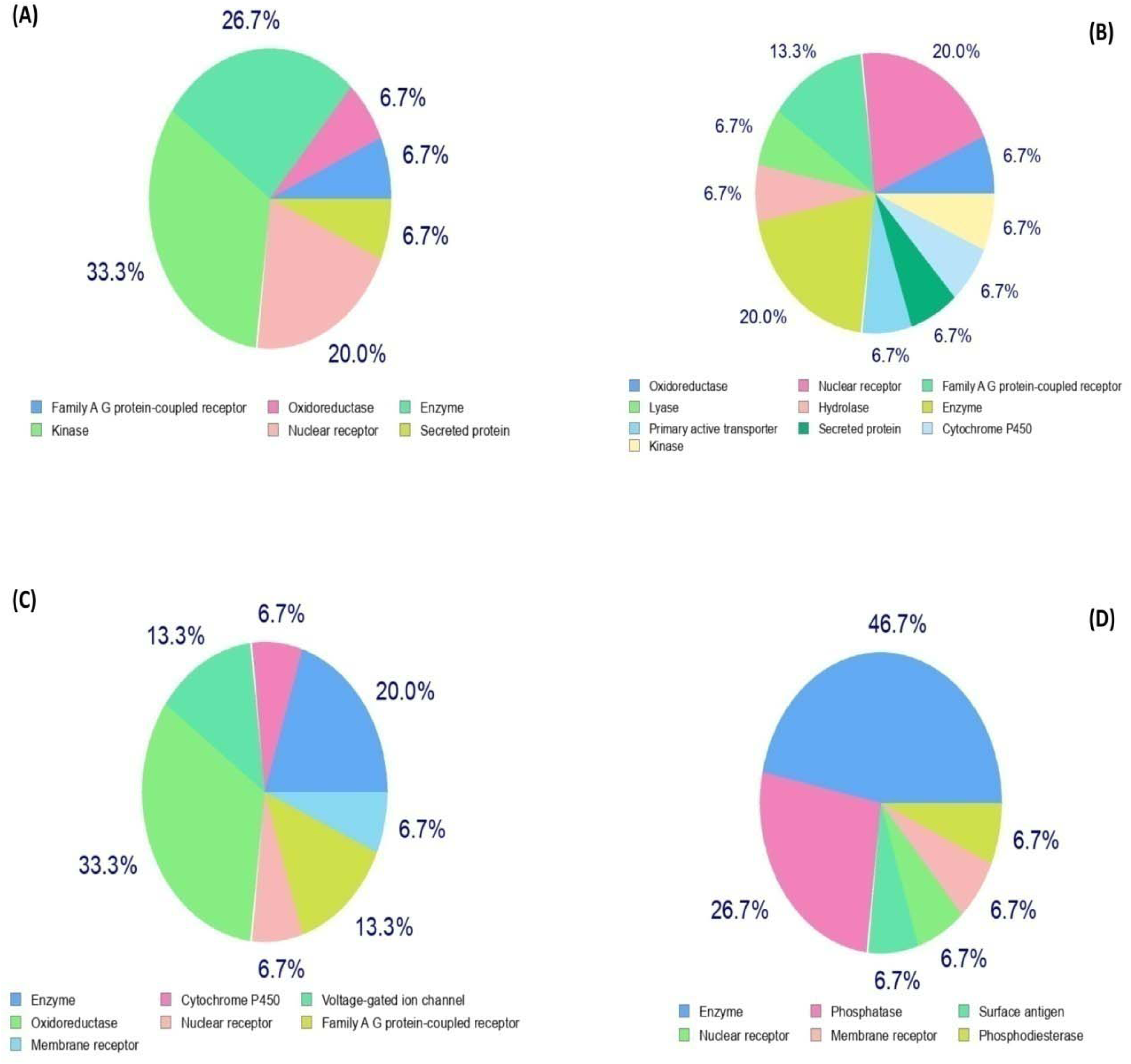
Predicted drug targets for gingerol (A), isoflavone (B), citronellal (C), asiatic acid (D).

**Figure 10:**
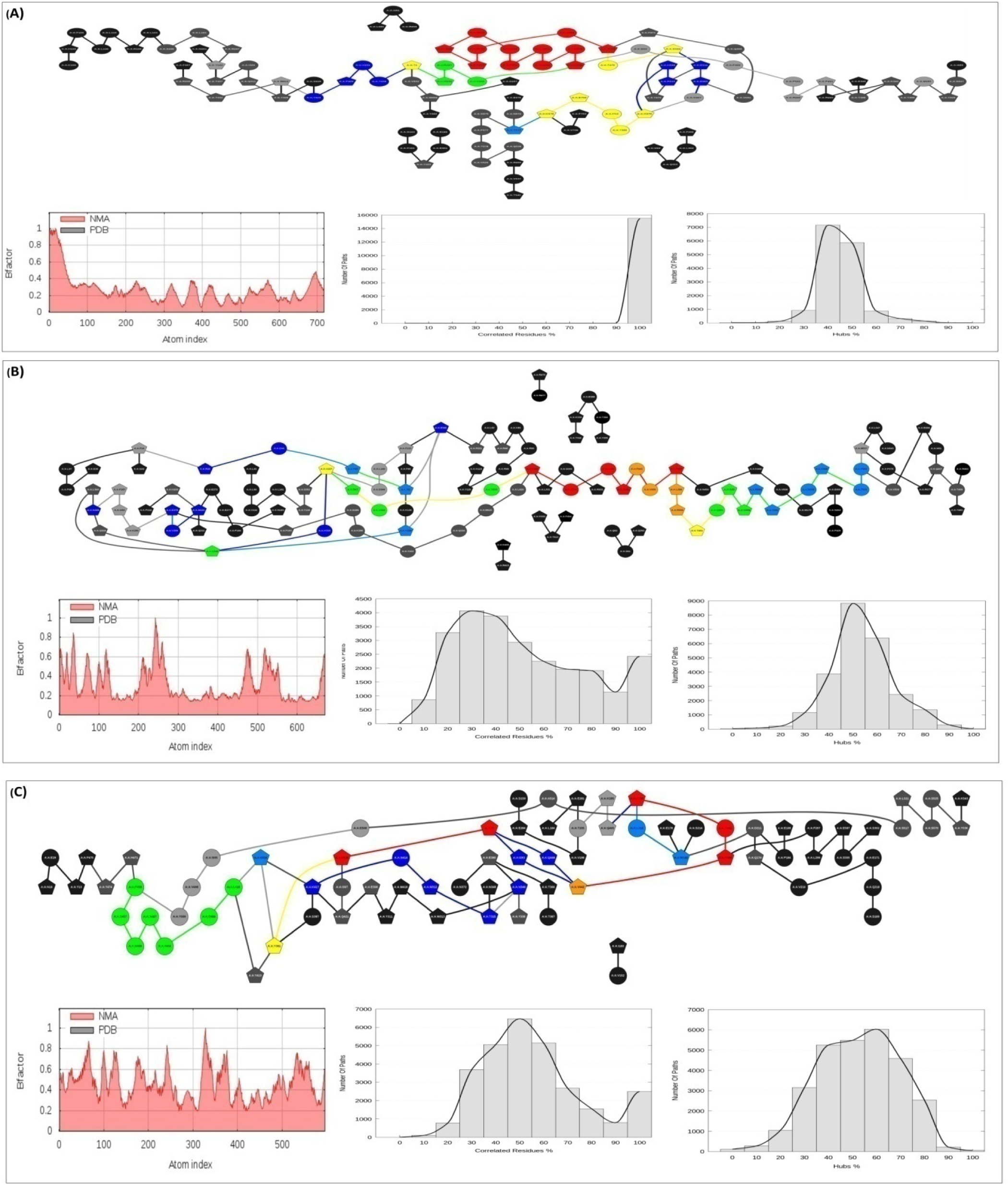
Metapathway, NMA, Corelated residues and Hub % analysis of MTA1-gingerol complex (A), MTA2-isoflavone omplex (B), and MTA3-isoflavone complex (C).

**Figure 11:**
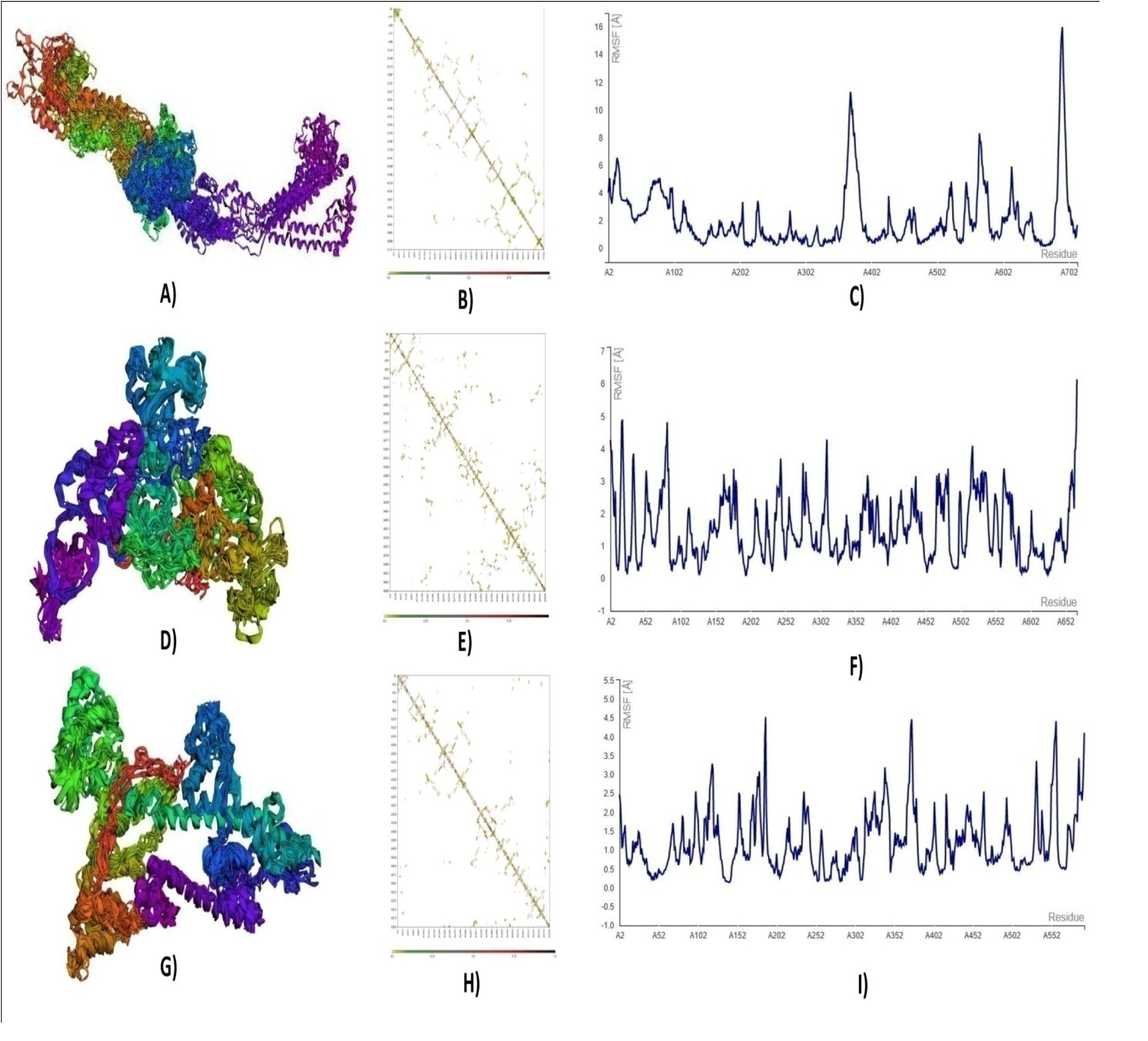
The final models (represented as cartoons) showed minor fluctuations for MTA1 (A), and comparatively rigidness for MTA2 (D) and MTA3 (G). The contact atom maps (B, E, H) and fluctuation plots of MTA1 (C), MTA2 (F) and MTA3 (I) represent the fluctuations in the residues during simulation.

### 3.8. Drug profile analysis of top metabolites

Several ADME properties including physicochemical parameters, pharmacokinetics, lipophilicity, water solubility and medicinal chemistry of top metabolites were demonstrated to evaluate their druggability potential (Figure 8). In this study, each drug candidate showed high GI absorption. Blood-brain barrier permeation computed by BOILED-Egg model revealed no BBB permeation for isoflavone, gingerol and asiatic acid. Besides, analysis of inhibition effects with different CYP isoforms (e.g. CYP1A2, CYP2D6, CYP2C9, CYP2C19, CYP3A4) confirmed that almost all candidates do not interact with cytochromes P450 isoforms. The metabolites showed bioavailability score of 0.5 and water solubility at moderate levels (Table 6).

**Table 6:**
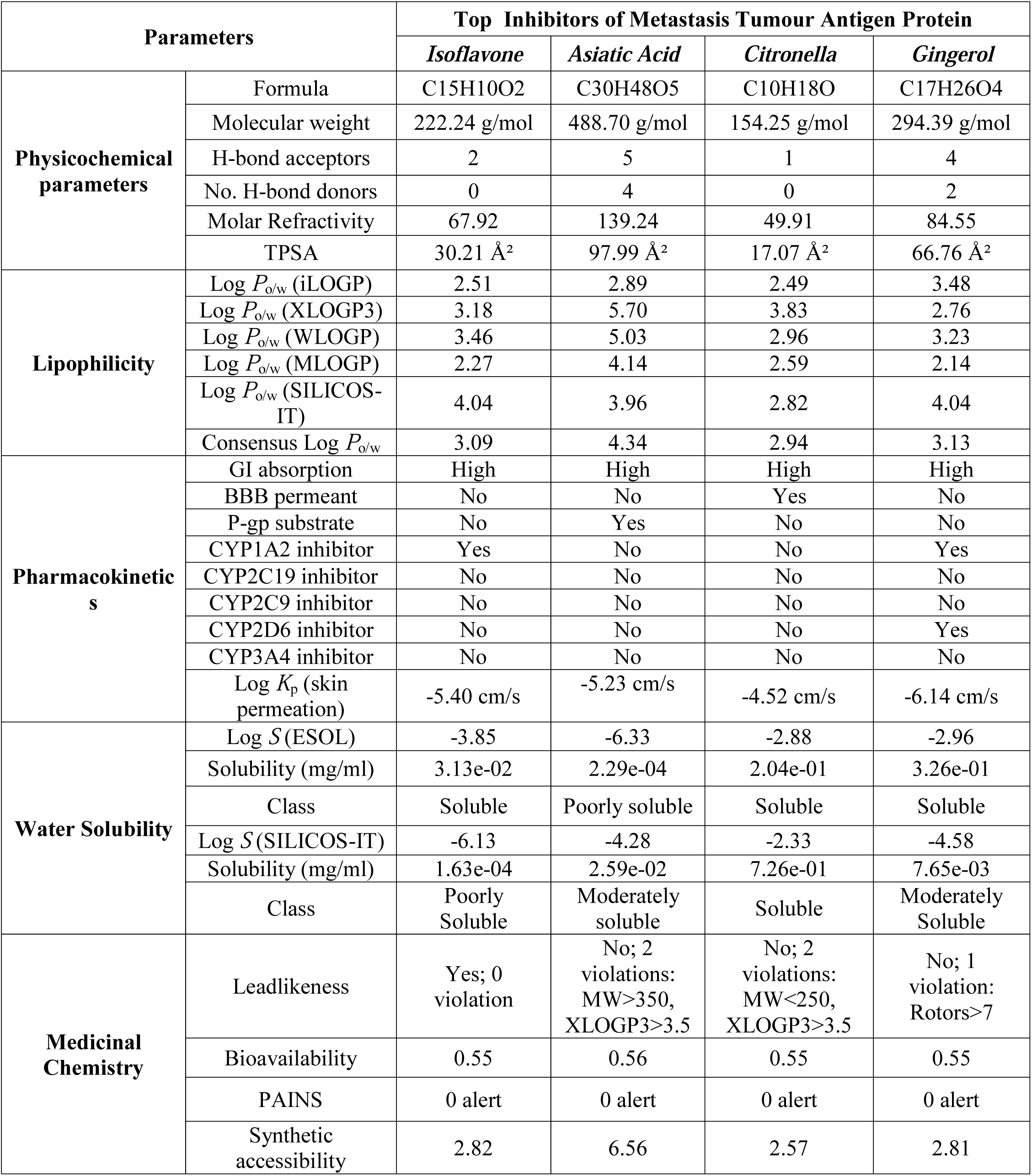
ADME analysis of top drug candidates

### 3.9. Toxicity pattern analysis of top metabolites

The relative toxicity (i.e AMES toxicity, oral rat toxicity, hepatotoxicity, skin sensitization, *T. pyriformis* toxicity, minnow toxicity)of top metabolites were predicted via pkCSM server. The results revealed negative outcomes in AMES test for isoflavone, gingerol, and asiatic acid indicating no risk of mutagenic or carcinogenic toxicity. The top drug candidates did not interact with hERGI (human ether-a-go-go related gene I) and hERG II,confirming that they are not hERG inhibitors. Moreover, top candidates revealed negative results for skin sensitization and hepatotoxicity. Minnow Toxicity values of all metabolites were greater than −0.3 log mM indicating them as non-toxic. Besides, maximum tolerated dose, oral rat acute toxicity (LD50), oral rat chronic toxicity (LOAEL) and *T. pyriformis* toxicity did not show any undesirable effects by the top drug candidates that could reduce their drug-likeness properties (Table 7).

**Table 7:** Toxicity pattern analysis of top drug candidates

| Toxicity Parameters | Top drug candidates |  |  |  |
| --- | --- | --- | --- | --- |
|  | Isoflavone | Gingerol | Citronellal | Asiatic Acid |
| AMES Test | No | No | Yes | No |
| Max. Tolerated Dose (log mg/kg/day) | 0.107 | 0.635 | 0.865 | 0.078 |
| hERG I inhibitor | No | No | No | No |
| hERG II inhibitors | No | No | No | No |
| Oral Rat Acute Toxicity, LD50 (mol/kg) | 1.726 | 1.958 | 2.482 | 2.592 |
| Oral Rat Chronic Toxicity, LOAEL (log mg/kg_bw/day) | 0.99 | 1.631 | 0.391 | 0.575 |
| Hepatotoxicity | No | No | No | No |
| Skin Sensitisation | No | No | No | No |
| *T. pyriformis* Toxicity (log µg/L) | 1.244 | 1.487 | 0.57 | 0.285 |
| Minnow Toxicity (log mM) | 0.603 | 0.966 | 4.555 | 1.106 |

### 3.10. Prediction of drug targets and available drug molecules from drugbank

Most of the target classes covered by the top drug candidates belonged to enzymes (e.g. oxidoreductase, phosphatase, lyase), nuclear receptors and secreted proteins (Figure 9, Table 8). SwissSimilarity prediction revealed that several approved drugs i.e. Tramadol, Estradiol, Nabumetone could be an alternative to Gingerol to block MTA proteins (Table 9). Moreover, Ethanolamine Oleate, Dihomo-gamma-linolenic acid (DGLA), 4-Androstenedione and some other synthetic analogs may be potent drugs which showed significant structural similarity with Citronella (Table 9).

**Table 8:**
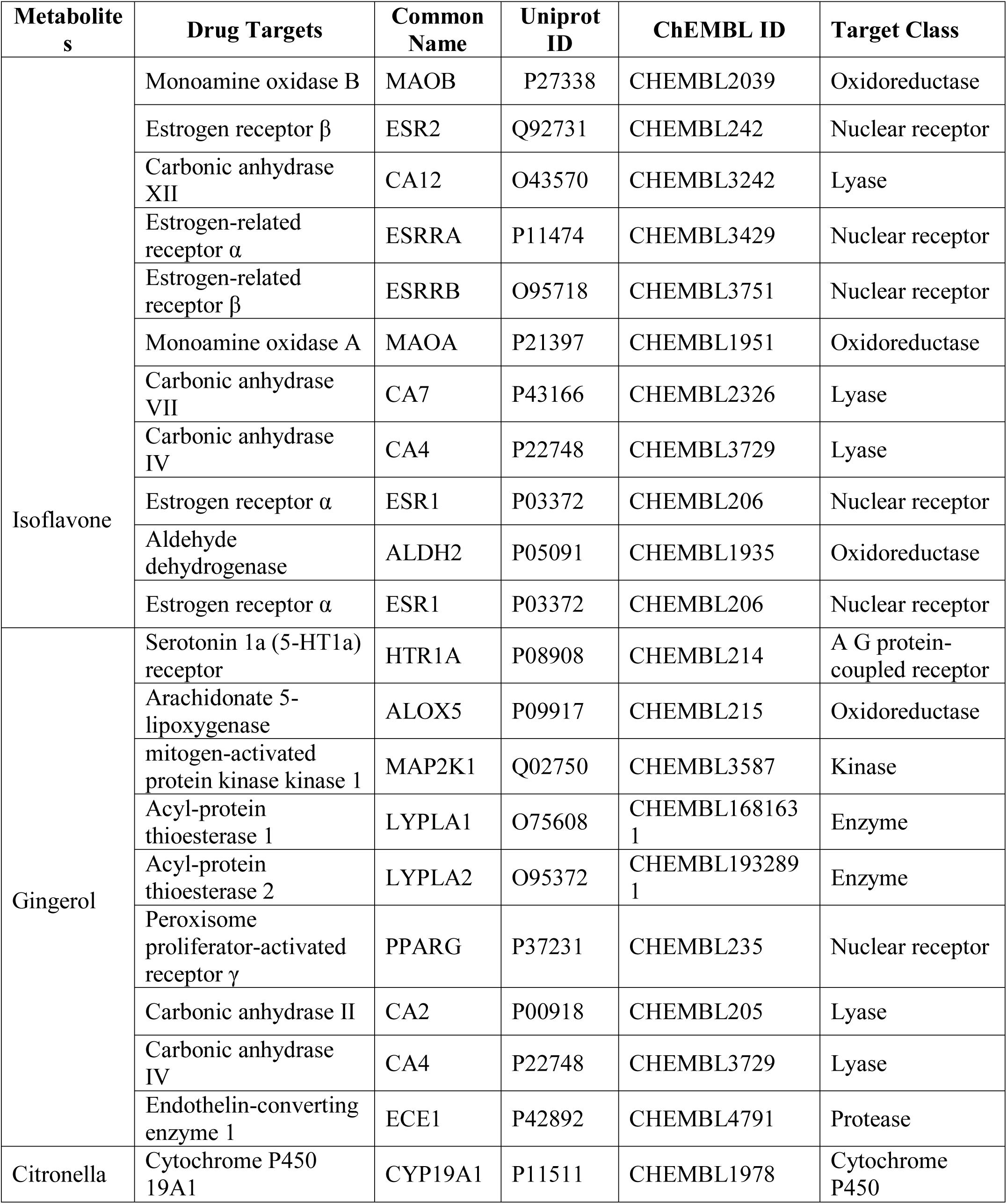
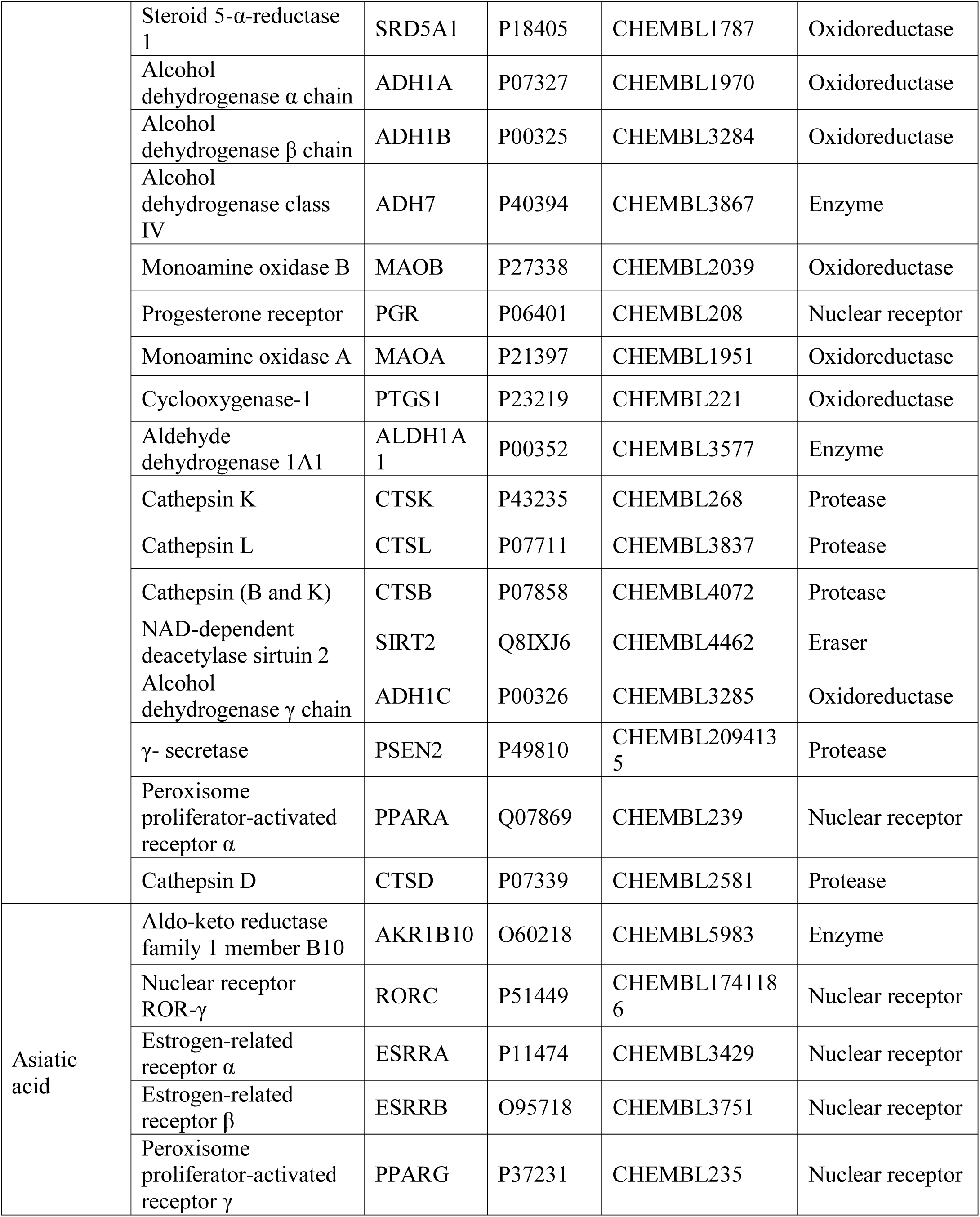
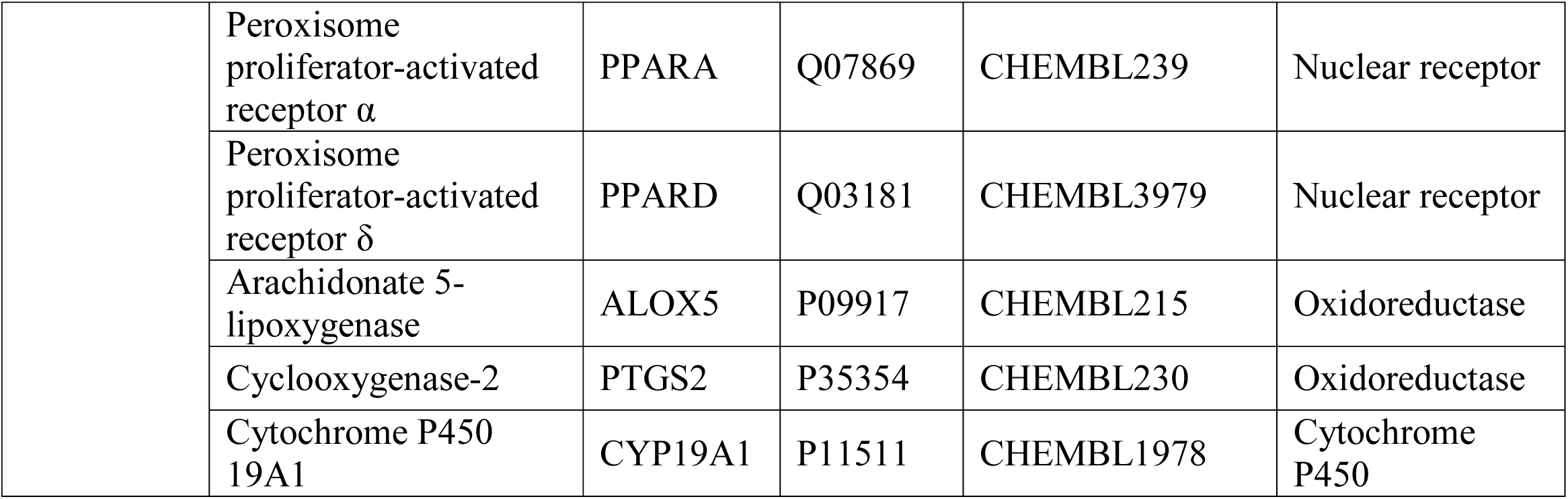
Predicted drug targets for Isoflavone, Gingerol, Citronellal and Asiatic acid

**Table 9:**
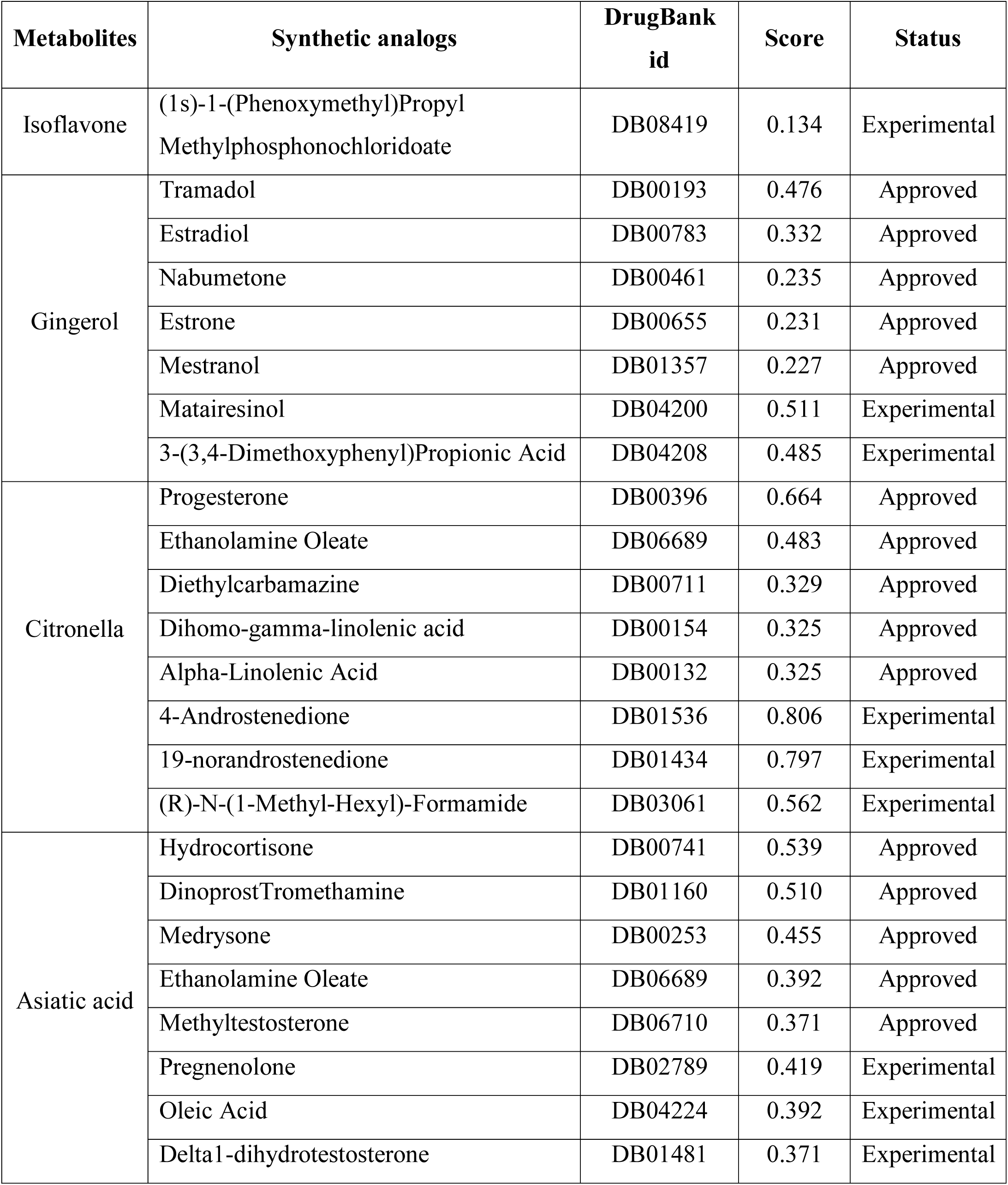
Predicted synthetic analogs of top metabolites from DrugBank

| Metabolites | Synthetic analogs | DrugBank id | Score | Status |
| --- | --- | --- | --- | --- |
| Isoflavone | (1s)-1-(Phenoxymethyl)Propyl Methylphosphonochloridoate | DB08419 | 0.134 | Experimental |
| Gingerol | Tramadol | DB00193 | 0.476 | Approved |
|  | Estradiol | DB00783 | 0.332 | Approved |
|  | Nabumetone | DB00461 | 0.235 | Approved |
|  | Estrone | DB00655 | 0.231 | Approved |
|  | Mestranol | DB01357 | 0.227 | Approved |
|  | Matairesinol | DB04200 | 0.511 | Experimental |
|  | 3-(3,4-Dimethoxyphenyl)Propionic Acid | DB04208 | 0.485 | Experimental |
| Citronella | Progesterone | DB00396 | 0.664 | Approved |
|  | Ethanolamine Oleate | DB06689 | 0.483 | Approved |
|  | Diethylcarbamazine | DB00711 | 0.329 | Approved |
|  | Dihomo-gamma-linolenic acid | DB00154 | 0.325 | Approved |
|  | Alpha-Linolenic Acid | DB00132 | 0.325 | Approved |
|  | 4-Androstenedione | DB01536 | 0.806 | Experimental |
|  | 19-norandrostenedione | DB01434 | 0.797 | Experimental |
|  | (R)-N-(1-Methyl-Hexyl)-Formamide | DB03061 | 0.562 | Experimental |
| Asiatic acid | Hydrocortisone | DB00741 | 0.539 | Approved |
|  | Dinoprost Tromethamine | DB01160 | 0.510 | Approved |
|  | Medrysone | DB00253 | 0.455 | Approved |
|  | Ethanolamine Oleate | DB06689 | 0.392 | Approved |
|  | Methyltestosterone | DB06710 | 0.371 | Approved |
|  | Pregnenolone | DB02789 | 0.419 | Experimental |
|  | Oleic Acid | DB04224 | 0.392 | Experimental |
|  | Delta1-dihydrotestosterone | DB01481 | 0.371 | Experimental |

## 4. Discussion

Despite extensive research and occasional successes, most of the cancer treatments are still a long way from reality [115]. Though treatments are available in some instances, not everyone can afford their prescriptions (one in five) and pay for treatments (one in four) [116]. Products of MTA gene family are involved and accelerate the migration, invasion, and survival of immortal cells in human. In this study, attempts were taken to reveal novel therapeutic option to prevent metastasis and cancer progression at early stages. Limited progress has been made so far in the treatment of cancer metastasis [117] and to the best of our knowledge similar computational strategies have not been yet explored to identify potential inhibitors of MTA gene products. The super-family, conserved domains and motifs of 3 MTA proteins i.e. MTA 1, MTA 2, MTA were analyzed to determine their functions. To date no crystal structure was determined for the MTA family proteins. Hence, homology modeling was performed for protein structure prediction from the available sequence data [118]. The 3D modeled structures were further studied extensively and validated through Ramachandran Plot and quality factor analysis (Figure 3-5). Moreover, ascertainment of stability can be done by comparing proteins essential dynamics to their normal modes [119, 120]. The refined models were significantly stable and showed minimum deformability at molecular level.

Virtual screening and de novo *drug* designing approaches are effective tools to identify novel hit molecules with desired biological activity [121, 122]. Analyzing the interaction between macromolecules and small ligands is an effective way to simplify the path of modern drug discovery [75], which also minimize the time and cost for drug development process [123]. In the next step, we performed structure based virtual screening for *in silico* evaluation of some plant based bioactive compounds as potent MTA inhibitors. Results showed that isoflavone, gingerol, citronellal and asiatic acid were the top hit molecules regarding minimum global binding energy. Asiatic acid, a triterpenoid derivative from *Centella asiatica*, was reported to have antioxidative, anti-angiogenic, anti-inflammatory, and neuro-protective activities [124, 125]. Both *in vitro* and *in vivo* studies confirmed the inhibition of lung and liver cancer by Asiatic acid [126, 127]. Citronella essential oils are often used as natural antimicrobials and showed to prevent the growth of harmful airborne bacteria [128, 129]. A recent study also revealed the potential use of citronella oil as chemotherapeutic agents against cancer [130]. Protective roles of plant flavonoids against cancer and heart disease were reported long ago [131]. Isoflavonoids were likely to be protective against a number of tumors and breast cancer [132]. In 2018, de Lima and colleagues reported the therapeutic potential of gingerol (*Zingiber officinale* extract) as anti cancer agent [133]. Previously gingerol showed to suppress colon cancer by targeting leukotriene A4 hydrolase [134]. Gingerol inhibits tumor growth and metastasis in human breast cells [135].

The drug surface hotspots and ligand binding interactions of the studied MTA proteins were unraveled (Table 5). Results showed that isoflovon and citronella occupied the BAH domains of MTA 1 and MTA 3 protein, respectively. In case of Asiatic acid, most of the ligand binding residues blocked the MTA-R1 domains of MTA 1 and MTA 2 protein (Table 5, Figure 7). The top drug candidates also inhibited the crucial binding sites of GATA and Myb-DNA binding domains of the MTA proteins. Most of the enzyme class (e.g. oxidoreductase, phosphatase, lyase) and nuclear receptors were identified as main targets by the top drug candidates (Table 8) carbonic anhydrases (CAs) have been shown to be important mediators of tumor cell pH by modulating the bicarbonate and proton concentrations for cell survival and proliferation [136]. This has led the researchers to inhibit specific CA isoforms, as an anti-cancer therapeutic strategy. Moreover, current evidence suggests that ECE-1c contributes to cancer aggressiveness and plays a putative role as a key regulator of cancer progression [137]. Alcohol/Aldehyde dehydrogenase (ADH) is used by cancer cells for energy, while the increase in total ADH in sera was positively correlated with renal *cancer* [138, 139]. Another enzyme γ*-secretase* is thought to play a significant *role* in tumorigenesis and pancreatic *cancer* [140]. The result suggests that all these enzymes can be specifically targeted by top drug candidates. Moreover, different nuclear receptors [141, 142], estrogen receptor beta (ERβ) [143], Dopamine D2 receptor [144], Progesteron receptor [145] are actively involved in metastasis and cancer progression. These receptors may also serve as effective pharmacological target by top metabolites (Table 5).

Optimization of ADME profiles is crucial for the clinical and commercial successes of drugs [146]. No side-effect was reported in some randomized, placebo-controlled trials of isoflavone and citronella essential oil [147, 148]. Therapeutic uses of gingerol and asiatic acid are also considered as safe and were supported by some pharmacokinetic and preclinical studies [149, 150]. In the present study, the ADME analysis did not find any undesirable consequences by the top drug candidates. However, cautions should be taken for the repurposed use of these drugs to avoid any adverse effects. The molecular dynamics studies have grown into a sophistication that enables macromolecular structural function relationships to be fully understood [151]. The deformability analysis of modelled proteins revealed that the distortions were lesser atomic distortion and also the higher eigenvalues of MTA1 (8.456619e^-08^), MTA2 (8.839058e^-05^) and MTA3 (1.015865e^-05^) are representative of higher energy which is required to deform the protein structures confirms the stability of modeled structures. Moreover, the docked complex B factor analysis revealed the lesser atomic distortions in the MTA1-gingerol and MTA2 -isoflavone docked complex but a bit in MTA3-isoflavone complex (Figure 10). The lesser atomic distortions also confirmed the stability of the complexes. The metapathway and hubs analysis shows interaction strength and cross-correlation of atomic motions of docked complex which implies to be a strong recurrent links between them (Figure 10). The minor fluctuations were observed in the RMSF plot, reflecting the uninterrupted interaction between the isoflavone and the MTA2, MTA3. Unlikely, higher fluctuations ranging over 10 **Å**, were observed in the RMSF plot, thus confirming a bit flexibility of the docked complex of gingerol and MTA1 (Figure 11).

We further performed ligand based virtual screening to identify similar drug molecules using a large collection of 3,76,342 compounds from DrugBank which are experimentally active on different macromolecular targets [112]. Results revealed that several synthetic analogs of gingerol (i.e. Tramadol, Estradiol, Nabumetone) may efficiently block the MTA proteins. Tramadol is a synthetic opioid which can be administered orally, subcutaneously, intravenously, intramuscularly, rectally and spinally [152]. It has been found to be a useful drug in patients with cancer pain (both with nociceptive and neuropathic characteristics) [153]. Estradiol is a major regulator of growth for the subset of breast cancers that express the estrogen receptor. Breast cancer cell lines that undergo long-term estrogen deprivation can be growth-inhibited by estradiol. High dose estrogen can be used to induce caspase-mediated death [154]. Nabumetone, on the other hand, can inhibit cyclooxygenase-COX enzyme and are thought to be tumor suppressor [155]. COX-2 and PG overexpression in chronic inflammation suggest that PGs produced by COX-2 catalysis can provide nutrition for tumor survival and proliferation [156]. Thus, Nabumetone can be used to inhibit COX and prostaglandin level which is a major factor for tumor formation. Results also suggest that Progesterone and Dihomo-gamma-linolenic acid (DGLA) may be an alternative choice to Citronella. Maintaining the balance of progesterone and unsaturated fatty acid have been found to exert clinical efficacy in arresting cancer cell and tumor growth [157, 158]. Another approved drug, Hydrocortisone showed high similarity with asiatic acid, which can be used for patients with advanced breast cancer. Though its mechanism of action is still unclear, a direct role in cell killing has been postulated for this glucocorticoid drug [159]. It is inevitable that computational target prediction provides valuable information for drug repurposing, understanding side effects and expanding the druggable genome [160-162]. Therefore, this study may pave the way of drug development against cancer metastasis in future.

## 5. Conclusion

The study suggests that isoflavone, gingerol, citronellal and asiatic acid could be potent MTA inhibitors to prevent cancer metastasis. Furthermore, several biologically active structural analogs from DrugBank i.e. Tramadol, Nabumetone, DGLA, Hydrocortisone may be effective and show potency to inhibit cancer progression in the early stage. However, the results are solely based on *in silico* investigations. Due to the encouraging results, we highly recommend further *in vitro* and *in vivo* trials using model animals for the experimental validation of the findings.

## Acknowledgements

Authors would like to acknowledge the Department of Microbial Biotechnology of Sylhet Agricultural University for the technical support of this research

