## Supplementary file 1 for "Identification of potential inhibitory analogs of metastasis tumor antigens (MTAs) using bioactive compounds: revealing therapeutic option to prevent malignancy"

Supplementary Table 2: Pocket ID, area, volume of MTA1 protein

| **Pocket ID** | **Area (SA)** | **Volume (SA)** | |
| --- | --- | --- | --- |
| 1 | 1318.734 | 557.492 | |
| 2 | 106.836 | 439.619 | |
| 3 | 185.960 | 284.723 | |
| 4 | 475.469 | 193.520 | |
| 5 | 368.242 | 166.498 | |
| 6 | 380.562 | 152.044 | |
| 7 | 77.727 | | 131.944 |
| 8 | 144.663 | | 115.961 |
| 9 | 134.493 | | 113.061 |
| 10 | 159.699 | | 65.422 |
| 11 | 155.055 | | 58.385 |
| 12 | 155.024 | | 54.859 |
| 13 | 125.954 | | 52.376 |
| 14 | 109.377 | | 49.707 |
| 15 | 26.672 | | 43.931 |
| 16 | 27.910 | | 40.128 |
| 17 | 42.807 | | 24.832 |
| 18 | 63.215 | | 23.653 |
| 19 | 25.764 | | 18.045 |
| 20 | 44.863 | | 17.966 |
| 21 | 81.891 | | 13.478 |
| 22 | 61.070 | | 10.375 |
| 23 | 46.838 | | 9.132 |
| 24 | 40.753 | | 8.708 |
| 25 | 23.534 | | 8.376 |
| 26 | 37.854 | | 7.489 |
| 27 | 29.413 | | 6.899 |
| 28 | 26.147 | | 6.669 |
| 29 | 21.789 | | 6.034 |
| 30 | 22.612 | | 5.548 |
| 31 | 15.877 | | 5.506 |
| 32 | 18.427 | | 4.103 |
| 33 | 12.566 | | 3.784 |
| 34 | 4.268 | | 3.417 |
| 35 | 16.409 | | 3.365 |
| 36 | 16.836 | | 3.358 |
| 37 | 5.765 | | 3.013 |
| 38 | 12.120 | | 2.987 |
| 39 | 7.807 | | 2.728 |
| 40 | 10.982 | | 2.714 |
| 41 | 14.263 | | 2.535 |
| 42 | 8.161 | | 1.717 |
| 43 | 4.029 | | 1.469 |
| 44 | 1.833 | | 1.318 |
| 45 | 8.942 | | 1.314 |
| 46 | 9.152 | | 1.123 |
| 47 | 12.668 | | 1.098 |
| 48 | 8.882 | | 1.072 |
| 49 | 11.714 | | 1.030 |
| 50 | 6.035 | | 0.951 |
| 51 | 5.464 | | 0.951 |
| 52 | 7.910 | | 0.849 |
| 53 | 5.418 | | 0.432 |
| 54 | 6.103 | | 0.400 |
| 55 | 6.014 | | 0.330 |
| 56 | 2.235 | | 0.286 |
| 57 | 2.576 | | 0.229 |
| 58 | 2.886 | | 0.221 |
| 59 | 1.308 | | 0.132 |
| 60 | 1.431 | | 0.125 |
| 61 | 2.495 | | 0.099 |
| 62 | 1.249 | | 0.085 |
| 63 | 1.998 | | 0.084 |
| 64 | 0.637 | | 0.073 |
| 65 | 1.470 | | 0.067 |
| 66 | 1.789 | | 0.066 |
| 67 | 1.107 | | 0.057 |
| 68 | 0.806 | | 0.054 |
| 69 | 1.091 | | 0.053 |
| 70 | 1.167 | | 0.051 |
| 71 | 0.849 | | 0.048 |
| 72 | 0.351 | | 0.045 |
| 73 | 1.007 | | 0.043 |
| 74 | 1.039 | | 0.042 |
| 75 | 0.835 | | 0.040 |
| 76 | -0.125 | | 0.039 |
| 77 | 0.477 | | 0.034 |
| 78 | 1.078 | | 0.032 |
| 79 | 0.696 | | 0.029 |
| 80 | 0.887 | | 0.029 |
| 81 | 0.686 | | 0.025 |
| 82 | 0.617 | | 0.024 |
| 83 | 0.681 | | 0.024 |
| 84 | -0.003 | | 0.019 |
| 85 | 0.472 | | 0.016 |
| 86 | 0.542 | | 0.015 |
| 87 | 0.441 | | 0.014 |
| 88 | 0.529 | | 0.014 |
| 89 | 0.435 | | 0.012 |
| 90 | 0.716 | | 0.011 |
| 91 | 0.464 | | 0.011 |
| 92 | 0.268 | | 0.008 |
| 93 | 0.278 | | 0.007 |
| 94 | 0.225 | | 0.006 |
| 95 | 0.347 | | 0.006 |
| 96 | 0.306 | | 0.006 |
| 97 | 0.172 | | 0.006 |
| 98 | 0.307 | | 0.005 |
| 99 | 0.246 | | 0.004 |
| 100 | 0.206 | | 0.004 |
| 101 | 0.145 | | 0.003 |
| 102 | 0.192 | | 0.003 |
| 103 | 0.123 | | 0.002 |
| 104 | 0.143 | | 0.002 |
| 105 | 0.115 | | 0.002 |
| 106 | 0.254 | | 0.002 |
| 107 | 0.149 | | 0.002 |
| 108 | 0.121 | | 0.002 |
| 109 | 0.127 | | 0.002 |
| 110 | 0.077 | | 0.001 |
| 111 | 0.080 | | 0.001 |
| 112 | 0.109 | | 0.001 |
| 113 | 0.098 | | 0.001 |
| 114 | 0.074 | | 0.001 |
| 115 | 0.057 | | 0.001 |
| 116 | 0.105 | | 0.001 |
| 117 | 0.016 | | 0.000 |
| 118 | 0.003 | | 0.000 |
| 119 | 0.010 | | 0.000 |
| 120 | 0.018 | | -0.000 |
| 121 | 0.047 | | 0.000 |
| 122 | 0.004 | | 0.000 |
| 123 | 0.003 | | 0.000 |
| 124 | 0.026 | | 0.000 |
| 125 | 0.001 | | 0.000 |
| 126 | 0.006 | | 0.000 |
| 127 | 0.032 | | 0.000 |
| 128 | 0.050 | | 0.000 |
| 129 | 0.007 | | 0.000 |
| 130 | 0.000 | | 0.000 |
| 131 | 0.000 | | 0.000 |
| 132 | 0.066 | | 0.000 |
| 133 | 0.001 | | 0.000 |
| 134 | 0.743 | | -0.086 |
| 135 | -1.118 | | -0.198 |

Supplementary Table 3: Pocket ID, area, volume of MTA 2 protein

| **Pocket ID** | **Area (SA)** | **Volume (SA)** |
| --- | --- | --- |
| 1 | 901.007 | 1563.224 |
| 2 | 1645.882 | 1516.759 |
| 3 | 1381.837 | 1222.693 |
| 4 | 854.980 | 1074.453 |
| 5 | 205.939 | 258.818 |
| 6 | 147.930 | 123.619 |
| 7 | 209.641 | 92.717 |
| 8 | 101.660 | 58.843 |
| 9 | 19.587 | 45.770 |
| 10 | 107.359 | 29.244 |
| 11 | 62.356 | 17.496 |
| 12 | 70.978 | 17.176 |
| 13 | 31.271 | 11.870 |
| 14 | 57.849 | 11.662 |
| 15 | 47.436 | 11.265 |
| 16 | 48.291 | 10.832 |
| 17 | 50.163 | 10.824 |
| 18 | 30.474 | 10.453 |
| 19 | 46.716 | 9.102 |
| 20 | 29.358 | 8.737 |
| 21 | 57.136 | 8.495 |
| 22 | 44.668 | 7.870 |
| 23 | 34.814 | 7.522 |
| 24 | 30.750 | 6.931 |
| 25 | 27.405 | 6.125 |
| 26 | 22.039 | 5.333 |
| 27 | 26.564 | 4.811 |
| 28 | 28.726 | 4.286 |
| 29 | 15.605 | 4.150 |
| 30 | 14.140 | 4.149 |
| 31 | 15.137 | 3.871 |
| 32 | 14.918 | 3.099 |
| 33 | 11.949 | 2.948 |
| 34 | 13.180 | 2.905 |
| 35 | 12.644 | 2.328 |
| 36 | 10.748 | 2.156 |
| 37 | 10.430 | 1.966 |
| 38 | 14.482 | 1.826 |
| 39 | 8.054 | 1.765 |
| 40 | 12.365 | 1.470 |
| 41 | 8.035 | 1.354 |
| 42 | 13.801 | 1.281 |
| 43 | 8.830 | 1.264 |
| 44 | 7.082 | 1.257 |
| 45 | 10.911 | 1.218 |
| 46 | 2.728 | 1.150 |
| 47 | 7.633 | 0.979 |
| 48 | 8.563 | 0.857 |
| 49 | 7.861 | 0.775 |
| 50 | 7.506 | 0.751 |
| 51 | 5.608 | 0.678 |
| 52 | 5.990 | 0.657 |
| 53 | 2.792 | 0.601 |
| 54 | 6.345 | 0.482 |
| 55 | 4.584 | 0.442 |
| 56 | 4.860 | 0.372 |
| 57 | 2.179 | 0.368 |
| 58 | 3.805 | 0.367 |
| 59 | 0.084 | 0.309 |
| 60 | 3.717 | 0.298 |
| 61 | 3.850 | 0.290 |
| 62 | 3.278 | 0.251 |
| 63 | 3.670 | 0.244 |
| 64 | 4.354 | 0.241 |
| 65 | 2.934 | 0.241 |
| 66 | 1.363 | 0.217 |
| 67 | 0.970 | 0.193 |
| 68 | 4.305 | 0.193 |
| 69 | 2.739 | 0.191 |
| 70 | 2.314 | 0.177 |
| 71 | 3.076 | 0.162 |
| 72 | 1.695 | 0.154 |
| 73 | 2.556 | 0.135 |
| 74 | 2.291 | 0.134 |
| 75 | 2.034 | 0.116 |
| 76 | 0.878 | 0.097 |
| 77 | 1.872 | 0.096 |
| 78 | 1.291 | 0.091 |
| 79 | 1.067 | 0.081 |
| 80 | 1.841 | 0.076 |
| 81 | 1.263 | 0.071 |
| 82 | 1.094 | 0.068 |
| 83 | 1.209 | 0.066 |
| 84 | 0.778 | 0.057 |
| 85 | 1.207 | 0.051 |
| 86 | 1.087 | 0.051 |
| 87 | 0.327 | 0.050 |
| 88 | 1.071 | 0.045 |
| 89 | 1.213 | 0.043 |
| 90 | 1.159 | 0.041 |
| 91 | 1.176 | 0.037 |
| 92 | 1.638 | 0.034 |
| 93 | 0.930 | 0.031 |
| 94 | 0.744 | 0.029 |
| 95 | 0.605 | 0.023 |
| 96 | 0.717 | 0.021 |
| 97 | 1.003 | 0.019 |
| 98 | 0.503 | 0.015 |
| 99 | 0.540 | 0.015 |
| 100 | 0.497 | 0.012 |
| 101 | 0.457 | 0.012 |
| 102 | 0.490 | 0.011 |
| 103 | 0.478 | 0.011 |
| 104 | 0.402 | 0.008 |
| 105 | 0.451 | 0.008 |
| 106 | 0.308 | 0.008 |
| 107 | 0.423 | 0.008 |
| 108 | 0.357 | 0.007 |
| 109 | 0.222 | 0.005 |
| 110 | 0.265 | 0.004 |
| 111 | 0.124 | 0.003 |
| 112 | 0.171 | 0.003 |
| 113 | 0.205 | 0.002 |
| 114 | 0.132 | 0.002 |
| 115 | 0.140 | 0.002 |
| 116 | 0.183 | 0.002 |
| 117 | 0.054 | 0.001 |
| 118 | 0.085 | 0.001 |
| 119 | 0.072 | 0.001 |
| 120 | 0.120 | 0.001 |
| 121 | 0.103 | 0.001 |
| 122 | 0.069 | 0.001 |
| 123 | 0.001 | 0.000 |
| 124 | 0.010 | 0.000 |
| 125 | 0.042 | 0.000 |
| 126 | 0.009 | 0.000 |
| 127 | 0.015 | 0.000 |
| 128 | 0.001 | 0.000 |
| 129 | 0.000 | 0.000 |
| 130 | 0.032 | 0.000 |
| 131 | 0.024 | 0.000 |
| 132 | 0.023 | 0.000 |
| 133 | 0.051 | 0.000 |
| 134 | 0.011 | 0.000 |
| 135 | 0.041 | 0.000 |
| 136 | 0.025 | 0.000 |
| 137 | 0.056 | 0.000 |
| 138 | 3.527 | -0.353 |

Supplementary Table 4: Pocket ID, area, volume of MTA 3 protein

| **Pocket ID** | **Area (SA)** | **Volume (SA)** |
| --- | --- | --- |
| 1 | 4094.143 | 3963.371 |
| 2 | 348.836 | 446.902 |
| 3 | 279.681 | 344.538 |
| 4 | 300.236 | 171.766 |
| 5 | 239.695 | 103.908 |
| 6 | 80.455 | 58.038 |
| 7 | 116.014 | 45.328 |
| 8 | 76.505 | 43.129 |
| 9 | 89.605 | 19.752 |
| 10 | 83.512 | 19.270 |
| 11 | 46.042 | 18.604 |
| 12 | 38.342 | 17.407 |
| 13 | 63.428 | 15.951 |
| 14 | 58.957 | 13.707 |
| 15 | 21.428 | 11.703 |
| 16 | 39.856 | 11.676 |
| 17 | 22.223 | 11.330 |
| 18 | 21.481 | 7.682 |
| 19 | 37.670 | 6.965 |
| 20 | 30.664 | 6.842 |
| 21 | 16.122 | 5.819 |
| 22 | 30.643 | 5.421 |
| 23 | 17.134 | 5.332 |
| 24 | 17.096 | 5.081 |
| 25 | 20.320 | 3.242 |
| 26 | 24.968 | 3.070 |
| 27 | 10.530 | 2.146 |
| 28 | 14.933 | 2.112 |
| 29 | 13.586 | 2.110 |
| 30 | 10.819 | 2.003 |
| 31 | 1.321 | 1.845 |
| 32 | 11.872 | 1.752 |
| 33 | 7.743 | 1.681 |
| 34 | 8.367 | 1.659 |
| 35 | 6.249 | 1.419 |
| 36 | 7.061 | 1.278 |
| 37 | 10.789 | 1.204 |
| 38 | 8.406 | 1.085 |
| 39 | 5.127 | 1.062 |
| 40 | 5.656 | 0.971 |
| 41 | 10.694 | 0.924 |
| 42 | 10.408 | 0.827 |
| 43 | 5.895 | 0.747 |
| 44 | 7.141 | 0.745 |
| 45 | 5.131 | 0.576 |
| 46 | 3.944 | 0.492 |
| 47 | 4.843 | 0.420 |
| 48 | 2.512 | 0.353 |
| 49 | 3.905 | 0.334 |
| 50 | 5.336 | 0.330 |
| 51 | 2.867 | 0.318 |
| 52 | 3.972 | 0.317 |
| 53 | 3.297 | 0.281 |
| 54 | 2.166 | 0.273 |
| 55 | 4.019 | 0.246 |
| 56 | 2.683 | 0.212 |
| 57 | 3.064 | 0.167 |
| 58 | 1.177 | 0.112 |
| 59 | 1.796 | 0.105 |
| 60 | 1.530 | 0.096 |
| 61 | 1.427 | 0.075 |
| 62 | 1.245 | 0.059 |
| 63 | 1.310 | 0.049 |
| 64 | 0.942 | 0.042 |
| 65 | 0.986 | 0.029 |
| 66 | 0.639 | 0.028 |
| 67 | 2.515 | 0.023 |
| 68 | 0.752 | 0.023 |
| 69 | 0.828 | 0.022 |
| 70 | 0.491 | 0.016 |
| 71 | 0.477 | 0.016 |
| 72 | 0.396 | 0.011 |
| 73 | 0.415 | 0.010 |
| 74 | 0.317 | 0.009 |
| 75 | 0.600 | 0.005 |
| 76 | 0.226 | 0.004 |
| 77 | 0.331 | 0.004 |
| 78 | 0.248 | 0.004 |
| 79 | 0.175 | 0.003 |
| 80 | 0.077 | 0.001 |
| 81 | 0.115 | 0.001 |
| 82 | 0.059 | 0.001 |
| 83 | 0.121 | 0.001 |
| 84 | 0.099 | 0.001 |
| 85 | 0.001 | 0.000 |
| 86 | 0.000 | 0.000 |
| 87 | 0.022 | 0.000 |
| 88 | 0.069 | 0.000 |
| 89 | 0.033 | 0.000 |
| 90 | 0.055 | 0.000 |
| 91 | 0.030 | 0.000 |
| 92 | 0.000 | 0.000 |
| 93 | 0.034 | 0.000 |
