## Supplementary file 2 for "Identification of potential inhibitory analogs of metastasis tumor antigens (MTAs) using bioactive compounds: revealing therapeutic option to prevent malignancy"

**Supplementary Table 5**: Docking result of 50 metabolites with MTA1, MTA2 and MTA3

| **Protein** | **Metabolites** | **Solution no.** | **Global energy** | **Attractive VdW** | **Repulsive VdW** | **ACE** | **HB** |
| --- | --- | --- | --- | --- | --- | --- | --- |
| MTA1 | Allicin | 6 | -31.01 | -9.48 | 0.79 | -10.95 | 0.00 |
|  | Allylpropyldisulfide | 9 | -29.68 | -12.16 | 2.33 | -8.37 | 0.00 |
|  | Andrographolide | 2 | -43.34 | -21.88 | 5.59 | -9.43 | 0.00 |
|  | Apigenin | 4 | -44.07 | -18.21 | 3.27 | -12.39 | 0.00 |
|  | Aristolochic acid |  | -42.88 | -26.88 | 6.77 | -12.93 | 0.00 |
|  | Ascorbic acid | 5 | -36.34 | -18.72 | 3.08 | -6.99 | 0.00 |
|  | Asiatic acid | 7 | -48.44 | -23.80 | 9.08 | -12.64 | 0.00 |
|  | Avicularin | 2 | -46.07 | -23.65 | 5.65 | -9.77 | 0.00 |
|  | Camphene | 9 | -28.93 | -12.64 | 1.87 | -7.29 | 0.00 |
|  | Capsaicin | 10 | -44.90 | -21.68 | 4.75 | -10.53 | 0.00 |
|  | Chavibetol | 1 | -28.79 | -12.01 | 0.22 | -6.94 | 0.00 |
|  | Chavicine | 1 | -44.27 | -22.30 | 8.02 | -10.83 | 0.00 |
|  | Chrysoeriol | 3 | -47.03 | -25.59 | 7.77 | -12.84 | 0.00 |
|  | Cinnamic | 1 | -28.62 | -11.98 | 0.82 | -7.17 | 0.00 |
|  | Citronella | 1 | -48.32 | -26.00 | 9.04 | -10.39 | 0.00 |
|  | Cianidanol | 2 | -29.25 | -9.82 | 3.51 | -10.85 | 0.00 |
|  | Cleomsclosin B | 5 | -47.01 | -21.38 | 4.74 | -11.94 | 0.00 |
|  | Coumadin | 1 | -45.15 | -20.92 | 7.66 | -12.66 | 0.00 |
|  | Curcumin | 3 | -42.62 | -23.16 | 8.37 | -9.89 | 0.00 |
|  | Dihydromorin | 7 | -36.42 | -17.50 | 3.30 | -8.02 | 0.00 |
|  | Eugenol | 4 | -24.98 | -11.82 | 0.56 | -4.83 | 0.00 |
|  | Flavylium | 2 | -48.11 | -23.21 | 2.46 | -9.61 | 0.00 |
|  | Galangin | 10 | -38.26 | -18.52 | 2.14 | -7.71 | 0.00 |
|  | Gentisic acid | 6 | -23.67 | -10.26 | 0.39 | -5.60 | 0.00 |
|  | Geraniol | 5 | -43.15 | -18.47 | 4.13 | -11.72 | 0.00 |
|  | Gingerol | 4 | -51.23 | -22.36 | 1.67 | -11.92 | 0.00 |
|  | Guajaverin | 1 | -43.91 | -25.70 | 8.71 | -8.25 | 0.00 |
|  | Hydrocyanic acid | 8 | -23.94 | -9.58 | 0.00 | -6.21 | 0.00 |
|  | Isoflavone | 10 | -41.78 |  |  | -8.06 | 0.00 |
|  | Kaempferol | 5 | -38.96 | -17.83 | 1.65 | -8.56 | 0.00 |
|  | Luteolin | 9 | -39.97 | -21.40 | 3.27 | -7.15 | 0.00 |
|  | Myrcene | 5 | -37.12 | -16.57 | 2.08 | -8.78 | 0.00 |
|  | Norartocarpeti | 7 | -40.27 | -18.84 | 3.08 | -9.13 | 0.00 |
|  | Paucine | 9 | -39.87 | -18.95 | 2.88 | -8.89 | 0.00 |
|  | Piperic acid | 1 | -33.35 | -15.79 | 2.00 | -7.26 | 0.00 |
|  | Piperine | 5 | -46.52 | -21.88 | 7.33 | -13.56 | 0.00 |
|  | procyanidin | 8 | -38.93 | -16.03 | 1.58 | -10.31 | 0.00 |
|  | Quercetine | 1 | -41.56 | -19.09 | 2.22 | -9.39 | 0.00 |
|  | Quinine | 4 | -44.07 | -18.21 | 3.27 | -12.39 | 0.00 |
|  | Reserpine | 3 | -16.16 | -10.82 | 0.00 | 1.23 | -1.79 |
|  | riboflavin | 10 | -29.43 | -12.93 | 1.28 | -6.97 | 0.00 |
|  | Steppogenin | 4 | -25.27 | -10.41 | 1.17 | -6.77 | 0.00 |
|  | Swertinin | 3 | -36.96 | -19.18 | 2.17 | -6.32 | 0.00 |
|  | Thymoquinone | 5 | -30.74 | -12.19 | 1.37 | -8.52 | 0.00 |
|  | Venilin | 2 | -46.89 | -22.32 | 0.82 | -8.95 | 0.00 |
|  | Vincamine | 7 | -45.08 | -22.10 | 4.38 | -10.26 | 0.00 |
|  | Vitexin | 9 | -45.78 | -25.60 | 4.92 | -13.46 | 0.00 |
|  | Withaferin | 4 | -48.51 | -22.04 | 10.00 | -15.14 | 0.00 |
|  | Yohimbine | 4 | -38.74 | -19.70 | 0.93 | -6.29 | 0.00 |
|  | Zingiberne | 5 | -35.44 | -16.32 | 1.66 | -7.78 | 0.00 |

|  | **Metabolites** | **Solution no.** | **Global energy** | **Attractive VdW** | **Repulsive VdW** | **ACE** | **HB** |
| --- | --- | --- | --- | --- | --- | --- | --- |
| MTA2 | Allicin | 1 | -29.67 | -9.65 | 1.37 | -10.26 | 0.00 |
|  | Allylpropyldisulfide | 8 | -29.67 | -12.64 | 1.11 | -7.34 | 0.00 |
|  | Andrographolide | 1 | -41.94 | -18.23 | 4.68 | -13.70 | 0.00 |
|  | Apigenin | 4 | -40.55 | -18.24 | 2.17 | -9.75 | 0.00 |
|  | Aristolochic acid | 9 | -46.45 | -20.19 | 2.21 | -11.74 | 0.00 |
|  | Ascorbic acid | 2 | -50.87 | -23.52 | 4.26 | -12.53 | 0.00 |
|  | Asiatic acid | 5 | -49.77 | -21.10 | 8.67 | -17.91 | 0.00 |
|  | Avicularin | 3 | -43.97 | -22.48 | 2.31 | -7.59 | 0.00 |
|  | Camphene | 1 | -28.96 | 11.28 | 0.90 | -8.24 | 0.00 |
|  | Capsaicin | 1 | -44.50 | -19.36 | 1.75 | -10.55 | 0.00 |
|  | Chavibetol | 3 | -32.43 | -12.49 | 0.29 | -8.80 | 0.00 |
|  | Chavicine | 1 | -48.95 | -20.68 | 2.37 | -12.53 | 0.00 |
|  | Chrysoeriol | 3 | -46.71 | -17.97 | 2.01 | -15.60 | 0.00 |
|  | Cinnamic | 2 | -29.03 | -10.96 | 1.64 | -8.74 | 0.00 |
|  | Citronellal | 7 | -44.09 | -16.99 | 2.98 | -15.35 | 0.00 |
|  | Cianidanol | 1 | -33.50 | -9.42 | 0.26 | -12.35 | 0.00 |
|  | Cleomsclosin B | 9 | -47.67 | -24.73 | 5.80 | -9.83 | 0.00 |
|  | Coumadin | 3 | -47.43 | -19.82 | 8.40 | -17.52 | 0.00 |
|  | Curcumin | 10 | -45.16 | -23.97 | 12.18 | -12.43 | 0.00 |
|  | Dihydromorin | 8 | -41.75 | -21.35 | 1.88 | -7.60 | 0.00 |
|  | Eugenol | 4 | -28.20 | -11.96 | 3.28 | -8.17 | 0.00 |
|  | Flavylium | 1 | -48.33 | -23.21 | 2.82 | -9.87 | 0.00 |
|  | Galangin | 3 | -43.57 | -19.61 | 5.85 | -12.13 | 0.00 |
|  | Gentisic acid | 4 | -30.33 | -11.94 | 0.02 | -7.80 | 0.00 |
|  | Geraniol | 7 | -46.99 | -19.53 | 4.15 | -13.19 | 0.00 |
|  | Gingerol | 3 | -52.71 | -24.66 | 7.39 | -13.83 | 0.00 |
|  | Guajaverin | 7 | -47.26 | -27.38 | 8.94 | -11.38 | 0.00 |
|  | Hydrocyanic acid | 2 | -28.54 | -12.89 | 2.85 | -7.32 | 0.00 |
|  | Isoflavone | 10 | -54.70 | -25.38 | 10.43 | -15.65 | 0.00 |
|  | Kaempferol | 4 | -45.58 | -20.39 | 1.37 | -10.40 | 0.00 |
|  | Luteolin | 4 | -46.18 | -18.45 | 0.96 | -12.39 | 0.00 |
|  | Myrcene | 7 | -38.78 | -14.60 | 1.58 | -11.51 | 0.00 |
|  | Norartocarpeti | 9 | -42.23 | -18.92 | 1.59 | -9.89 | 0.00 |
|  | Paucine | 2 | -51.58 | -20.48 | 2.65 | -14.64 | 0.00 |
|  | Piperic acid | 7 | -33.85 | -15.42 | 2.83 | -8.74 | 0.00 |
|  | Piperine | 10 | -43.47 | -21.28 | 11.49 | -13.65 | 0.00 |
|  | procyanidin | 9 | -38.96 | -17.58 | 4.14 | -10.17 | 0.00 |
|  | Quercetine | 3 | -45.42 | -20.86 | 2.59 | -10.57 | 0.00 |
|  | Quinine | 4 | -40.55 | -18.24 | 2.17 | -9.75 | 0.00 |
|  | Reserpine | 6 | -20.17 | -12.64 | 0.11 | 0.56 | -2.22 |
|  | riboflavin | 2 | -29.60 | -11.25 | 0.49 | -8.49 | 0.00 |
|  | Steppogenin | 2 | -25.64 | -10.64 | 0.98 | -6.63 | 0.00 |
|  | Swertinin | 1 | -41.14 | -14.99 | 0.91 | -12.27 | 0.00 |
|  | Thymoquinone | 5 | -27.87 | -11.48 | 0.16 | -6.85 | 0.00 |
|  | Venilin | 3 | -42.31 | -16.34 | 1.53 | -12.62 | 0.00 |
|  | Vincamine | 7 | -45.69 | -16.06 | 2.64 | -15.77 | 0.00 |
|  | Vitexin | 7 | -46.20 | -24.72 | 5.31 | -8.49 | 0.00 |
|  | Withaferin | 4 | -44.90 | -18.35 | 3.96 | -14.91 | 0.00 |
|  | Yohimbine | 3 | -47.46 | -19.63 | 0.01 | -11.35 | 0.00 |
|  | Zingiberne | 3 | -40.27 | -16.42 | 2.74 | -11.35 | 0.00 |

|  | **Metabolites** | **Solution no.** | **Global energy** | **Attractive VdW** | **Repulsive VdW** | **ACE** | **HB** |
| --- | --- | --- | --- | --- | --- | --- | --- |
| MTA3 | Allicin | 8 | -29.20 | -8.41 | 1.23 | -11.22 | 0.00 |
|  | Allylpropyldisulfide | 3 | -30.02 | -10.75 | 0.09 | -8.88 | 0.00 |
|  | Andrographolide | 3 | -41.12 | -20.23 | 7.50 | -11.00 | 0.00 |
|  | Apigenin | 5 | -44.77 | -18.23 | 1.06 | -11.54 | 0.00 |
|  | Aristolochic acid | 9 | -53.36 | -24.17 | 5.28 | -13.91 | 0.00 |
|  | Ascorbic acid | 8 | -44.67 | -18.94 | 0.10 | -10.42 | 0.00 |
|  | Asiatic acid | 4 | -53.22 | -24.66 | 7.16 | -14.20 | 0.00 |
|  | Avicularin | 8 | -45.70 | -22.99 | 6.43 | -10.46 | 0.00 |
|  | Camphene | 3 | -29.13 | -11.92 | 0.46 | -7.70 | 0.00 |
|  | Capsaicin | 3 | -42.90 | -19.65 | 2.52 | -9.84 | 0.00 |
|  | Chavibetol | 2 | -29.87 | -12.66 | 3.88 | -9.00 | 0.00 |
|  | Chavicine | 4 | -47.46 | -21.17 | 7.90 | -13.99 | 0.00 |
|  | Chrysoeriol | 1 | -53.17 | -23.95 | 8.96 | -16.03 | 0.00 |
|  | Cinnamic | 2 | -27.84 | -10.99 | 0.42 | -7.70 | 0.00 |
|  | Citronellal | 2 | -54.84 | -24.04 | 6.19 | -15.04 | 0.00 |
|  | Cianidanol | 4 | -31.79 | -9.75 | 0.00 | -10.96 | 0.00 |
|  | Cleomsclosin B | 2 | -48.01 | -19.38 | 6.20 | -15.08 | 0.00 |
|  | Coumadin | 3 | -52.14 | -26.79 | 14.41 | -14.95 | 0.00 |
|  | Curcumin | 4 | -49.57 | -23.88 | 1.38 | -9.68 | 0.00 |
|  | Dihydromorin | 5 | -39.61 | -18.04 | 2.50 | -9.14 | 0.00 |
|  | Eugenol | 1 | -30.47 | -11.91 | 0.26 | -8.31 | 0.00 |
|  | Flavylium | 2 | -54.08 | -24.56 | 4.50 | -13.41 | 0.00 |
|  | Galangin | 1 | -46.63 | -19.30 | 2.05 | -12.35 | 0.00 |
|  | Gentisic acid | 4 | -24.56 | -8.80 | 0.41 | -7.46 | 0.00 |
|  | Geraniol | 1 | -43.90 | -19.98 | 8.80 | -13.17 | 0.00 |
|  | Gingerol | 10 | -48.02 | -22.08 | 8.24 | -13.47 | 0.00 |
|  | Guajaverin | 10 | -52.43 | -23.66 | 4.20 | -12.91 | 0.00 |
|  | Hydrocyanic acid | 10 | -30.83 | -11.97 | 0.17 | -8.30 | 0.00 |
|  | Isoflavone | 9 | -69.20 | -29.52 | 6.22 | -18.74 | 0.00 |
|  | Kaempferol | 7 | -49.64 | -20.33 | 2.80 | -13.41 | 0.00 |
|  | Luteolin | 7 | -42.56 | -18.46 | 3.14 | -11.71 | 0.00 |
|  | Myrcene | 7 | -39.01 | -16.79 | 0.91 | -9.19 | 0.00 |
|  | Norartocarpeti | 9 | -39.20 | -17.29 | 3.70 | -10.39 | 0.00 |
|  | Paucine | 6 | -43.92 | -19.12 | 2.82 | -11.41 | 0.00 |
|  | Piperic acid | 1 | -36.24 | -16.01 | 0.25 | -7.90 | 0.00 |
|  | Piperine | 3 | -47.66 | -18.73 | 2.33 | -13.69 | 0.00 |
|  | procyanidin | 1 | -38.02 | -15.02 | 0.28 | -9.97 | 0.00 |
|  | Quercetine | 6 | -41.62 | -18.23 | 2.18 | -10.04 | 0.00 |
|  | Quinine | 5 | -44.77 | -18.23 | 1.06 | -11.54 | 0.00 |
|  | Reserpine | 1 | -15.57 | -10.95 | 1.02 | 2.40 | 0.00 |
|  | Riboflavin | 3 | -26.38 | -11.82 | 2.03 | -6.47 | 0.00 |
|  | Steppogenin | 4 | -24.05 | -10.09 | 0.61 | -5.91 | 0.00 |
|  | Swertinin | 1 | -41.82 | -18.98 | 1.94 | -9.39 | 0.00 |
|  | Thymoquinone | 3 | -30.49 | -12.99 | 1.08 | -7.43 | 0.00 |
|  | Venilin | 9 | -46.55 | -18.71 | 0.64 | -11.99 | 0.00 |
|  | Vincamine | 5 | -42.85 | -18.19 | 8.98 | -14.29 | 0.00 |
|  | Vitexin | 6 | -50.51 | -22.08 | 0.88 | -11.35 | 0.00 |
|  | Withaferin | 9 | -59.54 | -25.33 | 6.84 | -17.38 | 0.00 |
|  | Yohimbine | 1 | -46.01 | -18.83 | 2.11 | -12.23 | 0.00 |
|  | Zingiberne | 10 | -33.09 | -15.75 | 9.32 | -10.90 | 0.00 |
